# Copper is an essential regulator of the autophagic kinases ULK1/2 to drive lung adenocarcinoma

**DOI:** 10.1101/816587

**Authors:** Tiffany Tsang, Jessica M. Posimo, Andrea A. Gudiel, Michelle Cicchini, David M. Feldser, Donita C. Brady

**Affiliations:** Department of Cancer Biology, University of Pennsylvania, Philadelphia, PA, 19104, USA; Cell and Molecular Biology Graduate Group, University of Pennsylvania, Philadelphia, PA, 19104, USA; Abramson Family Cancer Research Institute, Perelman School of Medicine, University of Pennsylvania, Philadelphia, PA, 19104, USA

## Abstract

Targeting aberrant kinase activity in cancer relies on unmasking cellular inputs such as growth factors, nutrients, and metabolites that contribute to cancer initiation and progression^1^. While the transition metal copper (Cu) is an essential nutrient that is traditionally viewed as a static cofactor within enzyme active sites^2^, a newfound role for Cu as a modulator of kinase signaling is emerging^3, 4^. We discovered that Cu is required for the activity of the autophagic kinases ULK1/2 through a direct Cu-ULK1/2 interaction. Genetic loss of the Cu transporter *Ctr1* or mutations in ULK1 that disrupt Cu-binding reduced ULK1/2-dependent signaling and autophagosome complex formation. Elevated intracellular Cu levels are associated with starvation induced autophagy and sufficient to enhance ULK1 kinase activity and in turn autophagic flux. Targeting autophagy machinery is a promising therapeutic strategy in cancers^5^, but is limited by the absence of potent inhibitors and the emergence of resistance. The growth and survival of lung tumors driven by KRAS^G12D^ is diminished in the absence of *Ctr1*, depends on ULK1 Cu-binding, and is associated with reduced autophagy levels and signaling. These findings suggest a new molecular basis for exploiting Cu-chelation therapy to forestall autophagy signaling to limit proliferation and survival in cancer.

---

By default, the dynamic and adaptive nature of signaling networks allows them to respond and, in some cases, sense extracellular and intracellular inputs^6^. While growth factors, nutrients, and metabolites are well-appreciated regulators of cell proliferation, the contribution of transition metals to cellular processes that support proliferation and contribute to malignant transformation are understudied. The transition metal copper (Cu) is essential for a diverse array of biological processes from cellular proliferation, neuropeptide processing, free radical detoxification, and pigmentation^7^. The importance of intact Cu homeostatic mechanisms for cell growth control is underscored by the stunted growth and failure to thrive associated with Cu deficiency in Menkes disease patients^8^ and the increased prevalence of cancer in patients and animal models with hereditary Cu overload in Wilson disease^9–11^. However, the dysregulated growth phenotypes associated with alterations in Cu availability cannot be fully explained by the limited number of Cu-dependent enzymes that traditionally harness the redox potential of Cu as a structural or catalytic cofactor.

Several recent studies have illuminated novel functions for Cu as a non-structural intracellular mediator of signaling by directly connecting Cu to signaling pathway components that modulate cell proliferation or lipid metabolism^3, 4, 12^. We previously showed that in response to growth factors, Cu contributes to the amplitude of canonical MAPK signaling through an interaction between Cu and the kinases MEK1 and MEK2^3^. Examination of this new signaling mechanism in the context of cancer revealed that BRAF^V600E^-driven tumors depend on MEK1/2 association with Cu, and moreover, are sensitive to Cu-chelating drugs^4, 13, 14^. In Wilson disease, Cu influences 3’,5’-cyclic AMP (cAMP)-mediated lipolysis downstream of β-adrenergic receptor signaling by directly inhibiting the phosphodiesterase PDE3B^12^, serving as one molecular mechanism to explain the connection between aberrant Cu accumulation and altered lipid metabolism^10^. These findings bolster a new paradigm linking Cu levels to signaling pathway components that are essential for normal physiology, but that become dysregulated in multiple disease states including cancer. Thus, uncovering additional kinases that are regulated by Cu levels is needed to further define Cu-dependent cellular processes that may contribute to the efficacy of Cu chelators for cancer therapy^15^.

To explore other Cu-dependent kinases, we first performed an alignment of the Cu-binding sequence in MEK1^4^ against kinase domains and identified the autophagy regulatory kinases ULK1 and ULK2 as sharing significant sequence similarity at the amino acids (H188, M230, and H239) required for Cu-binding in MEK1 (Fig. 1a). Both ULK1 and ULK2 bind to a Cu-charged resin, but not a metal free resin control, Fe-, or Zn-charged resin (Fig. 1b,c). To investigate the impact of Cu-binding to ULK1 and ULK2 function, we first tested whether Cu is directly required for ULK1 and ULK2 kinase activity. ULK1 and ULK2 *in vitro* kinase activity was enhanced in response to increasing concentrations of CuCl_2_ (Fig. 1d,e and Supplementary Fig. 1e,f) and inhibited when treated with increasing concentrations of the Cu chelator tetrathiomolybdate (TTM) (Fig. 1f,g and Supplementary Fig. 1g,h). The reduced ULK1 and ULK2 kinase activity in presence of TTM was similar to inhibition achieved with the ULK1/2 inhibitor MRT68921^16^ (Fig. 1f,g and Supplementary Fig. 1g,h).

**Figure 1.**
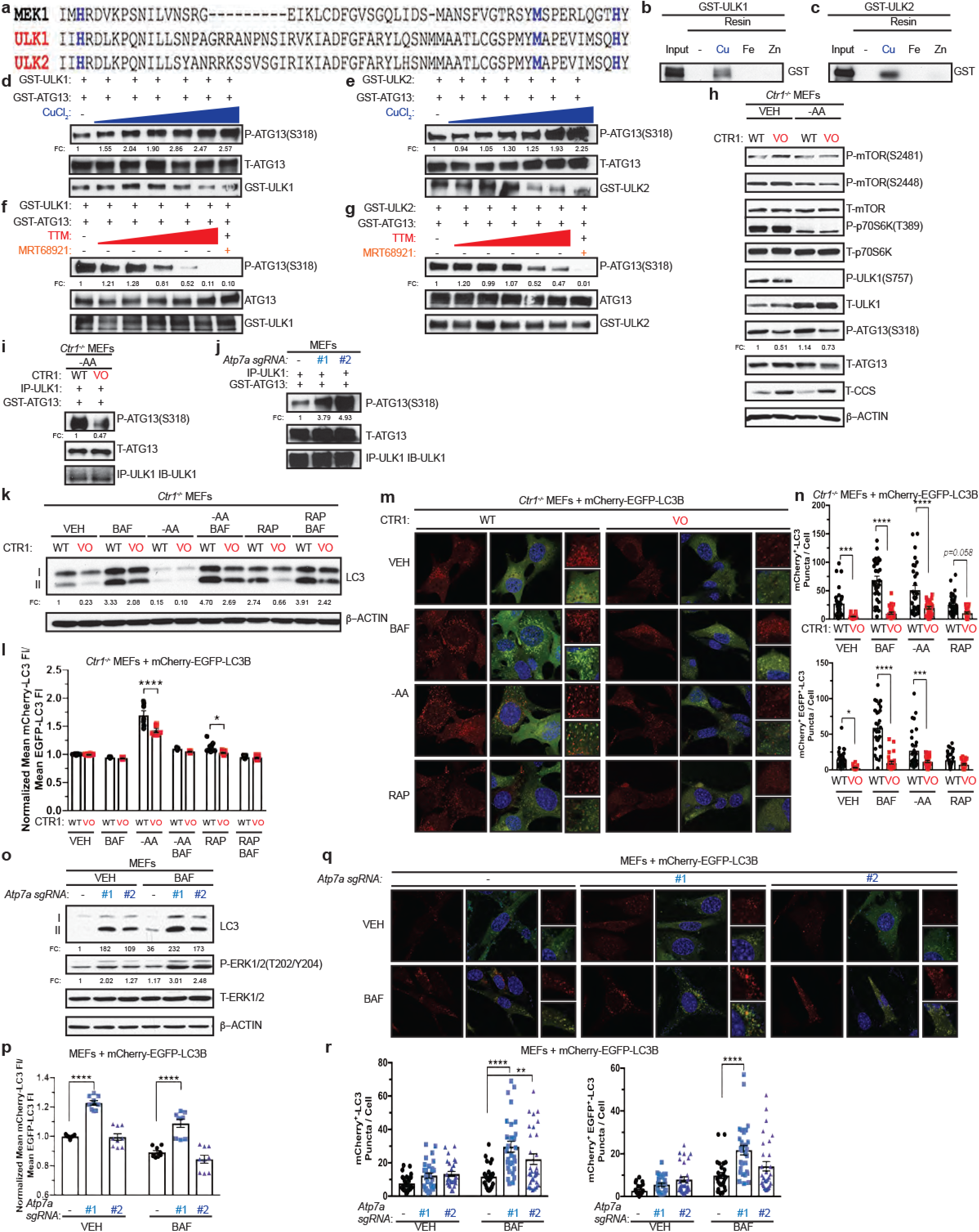
Cu binds to ULK1 and ULK2 and is required for kinase activity to promote autophagic flux. **a**, The amino-acid sequence of human MEK1 aligned to human ULK1 and human ULK2. Black letters, amino acids; blue letters, the four amino acids mutated in ***C***u-***b***inding ***m***utant (CBM) of MEK1 to decrease Cu binding and those conserved between ULK1 and ULK2. **b**,**c**, Immunoblot detection of recombinant GST-ULK1 or GST-ULK2 bound to a resin charged with or without Cu, Fe, or Zn compared to input. **d**,**e**,**f**,**g**, Immunoblot detection of recombinant phosphorylated (P)-ATG13, total (T)-ATG13, and GST-ULK1 or GST-ULK2 from ULK1 or ULK2 *in vitro* kinase assays treated with or without increasing concentrations of CuCl_2_ from 0 to 10 μM (**d**,**e**), with or without increasing concentrations of TTM from 0 to 50 μM, or 10 μM MRT68921 (**f**,**g**). Quantification: ΔP-ATG13/T-ATG13 normalized to GST-ULK1 and GST-ATG13 alone. **h**, Immunoblot detection of P- and/or T-mTOR, p70S6K, ULK1, ATG13, CCS, or β-ACTIN from *Ctr1^-/-^* MEFs stably expressing *CTR1^WT^* (WT) or *empty vector* (VO) treated with vehicle (VEH) or amino acid deprivation (-AA). Quantification: ΔP-ATG13/T-ATG13 normalized to WT, VEH control. **i**,**j**, Immunoblot detection of recombinant P-ATG13, T-ATG13, or immunoprecipitated (IP)-ULK1 from immunocomplex ULK1 kinase assays from *Ctr1^-/-^* MEFs stably expressing WT or VO treated with VEH or -AA or MEFs stably expressing *sgRNA* against *Rosa* (-) or *Atp7a* (*#1* or *#2*). Quantification: ΔP-ATG13/T-ATG13 normalized to WT, VEH control or *Rosa* (*-*) control. **k**, Immunoblot detection of LC3-I, LC3-II, or β-ACTIN from *Ctr1^-/-^* MEFs stably expressing WT or VO treated with VEH, -AA, or rapamycin (RAP), with or without bafilomycin (BAF). Quantification: ΔLC3-II/β-ACTIN normalized to WT, VEH control. **l**, Scatter dot plot of flow cytometry analysis of autophagic flux quantified by the ratio of mean mCherry-LC3 fluorescent intensity (FI)/mean EGFP-LC3 FI ± s.e.m. from *Ctr1^-/-^* MEFs stably expressing WT (black circles) or VO (red squares) and *mCherry-EGFP-LC3B* treated with VEH, -AA, or RAP, with or without BAF normalized to WT, VEH control. Results were compared using a two-way ANOVA followed by a Sidak’s multi-comparisons test. One asterisk, P<0.05; Four asterisks, P<0.0001. n≥7. **m**, Representative fluorescence images of EGFP, mCherry, or the merge from *Ctr1^-/-^* MEFs stably expressing WT or VO and *mCherry-EGFP-LC3B* treated with VEH, BAF, - AA, or RAP. **n**, Scatter dot plot of quantified mCherry positive (^+^)-LC3 puncta per cell or mCherry^+^ EGFP^+^-LC3 puncta per cell ± s.e.m. from *Ctr1^-/-^* MEFs stably expressing WT (black circles) or VO (red squares) and *mCherry-EGFP-LC3B* treated with VEH, BAF, -AA, or RAP. Results were compared using a two-way ANOVA followed by a Sidak’s multi-comparisons test. Three asterisks, P<0.001; two asterisks, P<0.0001. n≥24. **o**, Immunoblot detection of LC3-I, LC3-II, P-ERK1/2, T-ERK1/2, or β-ACTIN from MEFs stably expressing *sgRNA* against *Rosa* (-) or *Atp7a* (*#1* or *#2*) treated with VEH or BAF. Quantification: ΔLC3-II/T-β-ACTIN and ΔP-ERK1/2/T-ERK1/2 normalized to *Rosa* (*-*), VEH control. **p**, Scatter dot plot of flow cytometry analysis of autophagic flux quantified by the ratio of mean mCherry-LC3 FI/mean EGFP-LC3 FI ± s.e.m. from MEFs stably expressing *sgRNA* against *Rosa* (-, black circles) or *Atp7a* (*#1* or *#2*, blue squares, blue triangles) and *mCherry-EGFP-LC3B* treated with VEH or BAF normalized to *Rosa* (*-*), VEH control. Results were compared using a two-way ANOVA followed by a Sidak’s multi-comparisons test. Four asterisks, P<0.0001. n=9. **q**, Representative fluorescence images of EGFP, mCherry, or the merge from MEFs stably expressing *sgRNA* against *Rosa* (-) or *Atp7a* (*#1* or *#2*) and *mCherry-EGFP-LC3B* treated with VEH or BAF. **r**, Scatter dot plot of quantified mCherry^+^-LC3 puncta per cell or mCherry^+^ EGFP^+^-LC3 puncta per cell ± s.e.m. from MEFs stably expressing *sgRNA* against *Rosa* (-, black circles) or *Atp7a* (*#1* or *#2*, blue squares, blue triangles) and *mCherry-EGFP-LC3B* treated with VEH or BAF. Results were compared using a two-way ANOVA followed by a Sidak’s multi-comparisons test. Two asterisks, P<0.01; Four asterisks, P<0.0001. n≥26. Western blots images are representative of at least three biological replicates.

ULK1/2 are well-established downstream targets of the protein kinase mammalian target of rapamycin (mTOR)^17–20^. When associated with rapamycin-sensitive mTOR complex 1 (mTORC1)^21^, mTOR controls cellular energetics by driving anabolic processes, such as lipid, nucleotide, and protein synthesis^22^, and suppressing catabolic processes^17–20^, like autophagy. mTOR kinase activity is blocked in response to amino acid deprivation^23–25^, low cellular ATP levels^26, 27^, or rapamycin treatment^28, 29^. This in turn relieves the inhibitory phosphorylation of ULK1 and ULK2 and initiates the formation of an autophagosome^17–20^, which degrades and recycles proteins, lipids, and organelles to restore nutrient balance in the form of building blocks for metabolic pathways. To investigate the relationship between Cu and mTORC1-ULK1/2 signaling in mammalian cells, Cu-replete and Cu-deficient cells were generated by expressing the high affinity Cu transporter CTR1 (WT) or an empty vector (VO) in immortalized *Ctr1^-/-^* mouse embryonic fibroblasts (MEFs). Intracellular Cu levels in CTR1^WT^ expressing *Ctr1*-null MEFs are comparable to *Ctr1^+/+^* MEFs as confirmed by reduced CCS protein stability^30^, which is degraded in a Cu-dependent fashion, increased Cu levels via inductively coupled plasma mass spectrometry (ICP-MS), and elevated fluorescence from the CF4 probe^31^, which detects labile Cu (Fig. 1d and Supplementary Fig. 1a-d). *Ctr1* loss reduced ULK1-mediated phosphorylation of ATG13 under basal conditions and the elevated phosphorylation of ATG13 that occurs upon amino acid deprivation^17–20^ was only observed in the Cu-replete conditions (Fig. 1d and Supplementary Fig. 1i). In contrast, PI3K/AKT signaling-mediated phosphorylation^32^ of mTOR^S2448^ and mTOR phosphorylation^33–35^ of ULK1^S757^, p70S6K^T389^, and mTOR^S2481^ remained unchanged upon reduced Cu levels (Fig. 1d). These data suggest that Cu is required for ULK1 activity but not its upstream regulation by mTORC1 or mTOR itself. In agreement, ULK1 kinase activity is elevated upon amino acid withdrawal in Cu-replete cells but is blunted in the Cu-deficient cells (Fig. 1i and Supplementary Fig. 1j). Additionally, MEFs lacking the Cu exporter *Atp7a*, which have elevated intracellular Cu levels, also had increased ULK1 kinase activity (Fig. 1j and Supplementary Fig. 1k). These findings that reduced Cu levels decrease ULK1/2-dependent signaling downstream of mTORC1 inhibition (Fig. 1h-j) suggest that autophagy may be a newly identified Cu-dependent process.

The ULK1-ATG13 complex senses low-energy and nutrient depleted cellular states and initiates autophagosome formation^17–20^. To determine whether changes in levels of Cu affect ULK1/2-dependent autophagy, Cu-replete or Cu-deficient MEFs were exposed to amino acid deprivation^23–25^ or rapamycin^28, 29^, which inhibit mTOR. Specifically, autophagic flux was measured by three techniques, including *i*) Western blot analysis of the processing of LC3 from its full-length form (LC3-I) to its cleaved and phosphatidylethanolamine-conjugated form (LC3-II)^36^, *ii*) flow cytometry measurement of the ratio of autolysosomes (mCherry-LC3B) to autophagosomes (EGFP-LC3B)^37^, and *iii)* immunofluorescence detection of the abundance of autophagosomes (mCherry-EGFP-LC3 positive puncta) and autolysosomes (mCherry-LC3 positive puncta)^38^. These studies revealed that loss of *Ctr1* significantly reduced LC3-I processing (Fig. 1k and Supplementary Fig. 1l) and prevented autophagosome to autolysosome conversion upon autophagy induction (Fig. 1l-n). Further, under basal conditions the number of mCherry-EGFP-LC3 positive puncta was significantly reduced, indicating a potential block in autophagosome formation and in turn recruitment of LC3B (Fig. 1m,n). In contrast, CRISPR/Cas9-mediated knockdown of endogenous *Atp7a* (Supplementary Fig. 1m) caused elevated Cu levels that were sufficient to enhance LC3-II levels (Fig. 1o) and autophagic flux (Fig. 1p-r). Collectively, these data suggest that the levels of Cu could serve as a rheostat for ULK1/2-dependent autophagy by regulating ULK1/2 kinase activity in an mTOR-dependent fashion.

Since altering Cu levels has been shown to affect MAPK signaling^3, 4^, we interrogated the contribution of the MAPK pathway to Cu-mediated regulation of autophagy via genetic and pharmacologic approaches. First, we directly tested whether disrupting MEK1 Cu-binding was sufficient to reduce autophagy in the presence (*Ctr1^flox/flox^*) or absence (*Ctr1^-/-^*) of Cu transport. While expression of ***C***u-***b***inding ***m***utant (CBM) of MEK1^4^ reduced phosphorylation of ERK1/2 (Supplementary Fig. 1o), LC3-II levels were unchanged and Cu-dependent autophagic flux was still observed in *Ctr1^-/-^* MEFs (Supplementary Fig. 1p,q). Second, treatment with the MEK1/2 inhibitor, trametinib, which abolished phosphorylation of ERK1/2, did not phenocopy the reduced autophagic flux in the absence of *Ctr1* (Supplementary Fig. 1r,s). Consistent with these findings, expression of a gain-of-function (GOF) ERK2 mutant^4^, which bypasses the ability of Cu to influence MEK1/2 activity in *Ctr1^-/-^* MEFs (Supplementary Fig. 1t), did not increase LC3-II levels in cells devoid of significant Cu transport (Supplementary Fig. 1u,v). Finally, while increasing Cu levels via *Atp7a* knockdown is sufficient to increase phosphorylation of ERK1/2 (Fig. 1o), loss of *Atp7a* in MEFs lacking the key autophagy regulator *Atg5* (Supplementary Fig. 1m), which is part of the ubiquitin-like conjugation system that lipidates LC3^36^, failed to re-establish autophagy (Supplementary Fig. 1w). Collectively, these data indicate that Cu-dependent autophagy is mechanistically independent of Cu-regulated MEK1/2 activity and upstream of ATG5, which is required for autophagosome nucleation.

Since our findings suggest that Cu is required for ULK1/2-dependent signaling and autophagic flux in response to mTOR inhibition, we next hypothesized that Cu levels may fluctuate in an acute manner after autophagy induction. Interestingly, a statistically significant elevation in labile intracellular Cu levels, as measured by increased fluorescence of the Cu probe CF4^31^ and not its control (Ctrl-CF4)^31^, was detected in a time-dependent fashion post amino acid withdrawal (Fig. 2a,b, Supplementary Fig. 2a,b, and Supplementary Video 1-4). Thus, the requirement for Cu for efficient autophagic flux is driven by Cu fluctuations that support ULK1/2 kinase activity. While ULK1/2 activation is typically a prerequisite for LC3 processing by the ubiquitin-like conjugation system on the forming autophagosome^17^, several upstream ULK1/2-dependent molecular events are essential to initiate autophagosome formation^39, 40^ (Supplementary Fig. 2c). Specifically, active ULK1 translocates to the ER with its constitutive binding partners ATG13 and FIP200 to drive phagophore formation by phosphorylating core components of the class III PI3K VPS34 complex, which includes BECLIN-1, PIK3R4 (p150), and VPS34 itself^17–20, 41–43^. The ULK1-mediated phosphorylation of BECLIN-1 or ATG14 within VPS34 complex 1 stimulates VPS34 activity, which converts phosphatidylinositol to PI3P^43, 44^. The generation of PI3P serves as a docking site for FYVE domain containing proteins, like DFCP1 and WIPI1, that recruit the ubiquitin like conjugation complex ATG5-ATG12-ATG16 to lipidate LC3B^39, 45^. To dissect the contribution of Cu to the molecular events directly downstream of ULK1-dependent autophagosome formation, we visualized ULK1 complex translocation to puncta, (EGFP-ATG13 or EGFP-FIP200), VPS34 complex PI3P generation on puncta (EGFP-DFCP1), and PI3P effector translocation to puncta (EGFP-WIPI1) in response to amino acid deprivation in the presence (*Ctr1^flox/flox^*) or absence (*Ctr1^-/-^*) of Cu transport. Cu deficiency significantly decreased both ATG13, FIP200, WIPI1, and DFCP1 puncta formation in response to amino acid deprivation, indicating that Cu transport contributes to the early steps of phagophore nucleation (Fig. 2c-j). Defective recruitment of these initiating complexes and PI3P production, limited the number of EGFP-LC3 positive autophagosomes measured by flow cytometry^46^ (Fig. 2k and Supplementary Fig. 2d), an effect that was rescued by loss of the Cu exporter *Atp7A* (Fig. 2l). A selective requirement for Cu in the regulation of autophagosome formation was observed, as the Cu chelator TTM reduced EGFP-LC3 positive autophagosome number, while the iron (Fe) chelator desferoxamine (DFO) had no effect (Supplementary Fig. 2e,f). These findings suggest that ULK1/2-driven autophagy initiation requires Cu transport.

**Figure 2.**
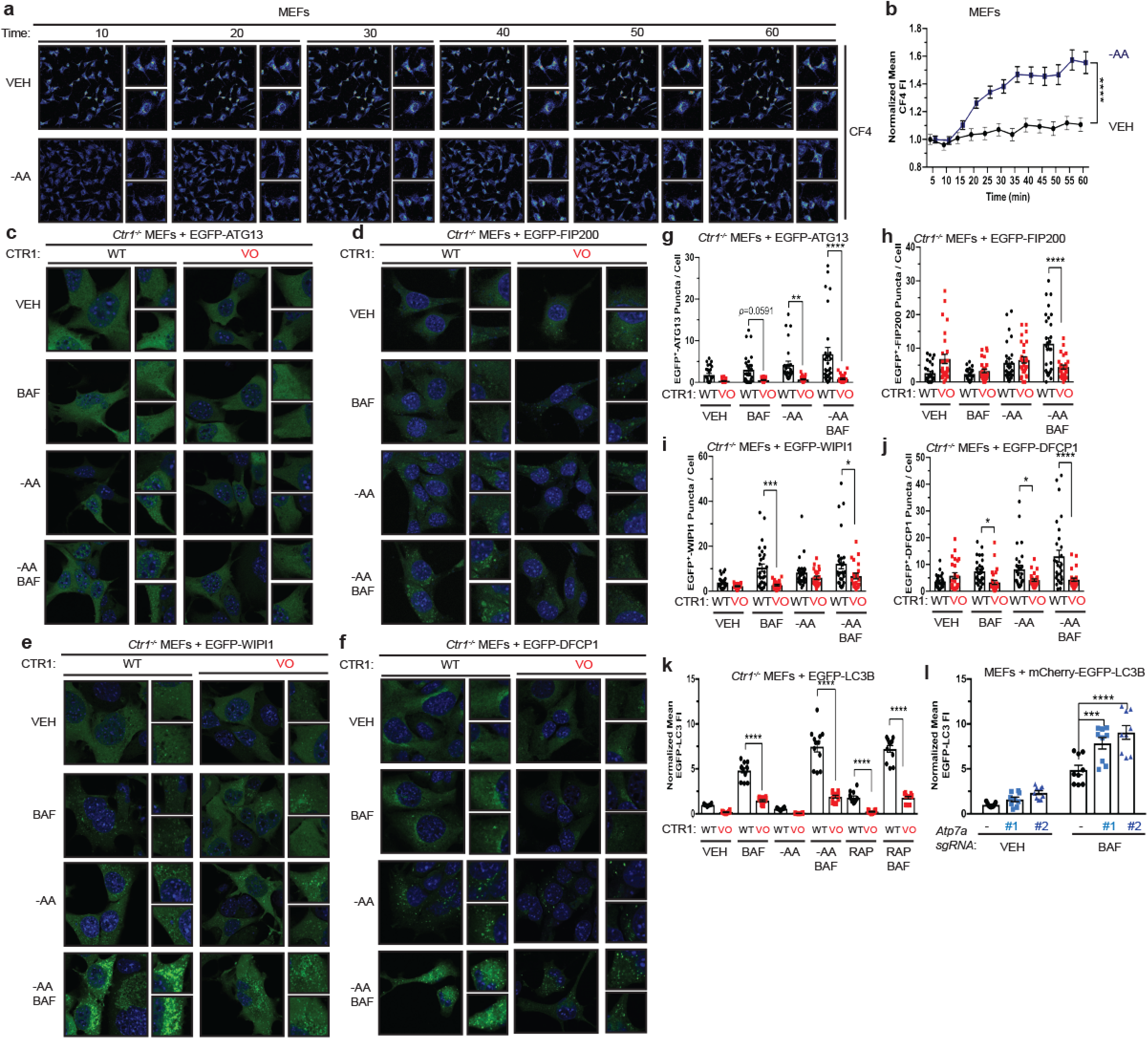
Cu is both necessary and sufficient for autophagosome formation. **a**, Representative live cell imaging of the Cu probe CF4 every ten minutes for 60 minutes from MEFs treated with vehicle (VEH) or amino acid deprivation (-AA). **b**, Quantification of mean CF4 fluorescence intensity (FI) ± s.e.m. versus time (minutes, min) from MEFs treated with VEH (black circles) or -AA (blue squares) normalized to t=0, five minutes. Results were compared using a two-way ANOVA followed by a Sidak’s multi-comparisons test. Four asterisks, P<0.0001. n=30. **c**,**d**,**e**,**f**, Representative fluorescence images of EGFP-ATG13 (**c**), EGFP-FIP200 (**d**), EGFP-WIP1 (**e**), or EGFP-DFCP1 (**f**) from *Ctr1^-/-^* MEFs stably expressing *CTR1^WT^* (WT) or *empty vector* (VO) and *EGFP-ATG13*, *EGFP-FIP200*, *EGFP-WIP1*, or *EGFP-DFCP1* treated with VEH or -AA with or without bafilomycin (BAF). **g**,**h**,**i**,**j**, Scatter dot plot of quantified EGFP^+^ puncta per cell ± s.e.m. from *Ctr1^-/-^* MEFs stably expressing WT (black circles) or VO (red squares) and *EGFP-ATG13* (**g**), *EGFP-FIP200* (**h**), *EGFP-WIP1* (**i**), or *EGFP-DFCP1* (**j**) treated with VEH or -AA with or without bafilomycin (BAF). Results were compared using a two-way ANOVA followed by a Sidak’s multi-comparisons test. One asterisk, P<0.05; Two asterisks, P<0.01; Three asterisks, P<0.001; Four asterisks, P<0.0001. EGFP-ATG13, ≥26; EGFP-FIP200, n≥21; EGFP-WIP1, n≥18; EGFP-DFCP1, n≥24. **k**, Scatter dot plot of flow cytometry analysis of the number of LC3-II positive autophagosomes quantified by the mean EGFP-LC3 FI ± s.e.m. from *Ctr1^-/-^* MEFs stably expressing WT (black circles) or VO (red squares) and *EGFP-LC3B* treated with VEH, -AA, or RAP with or without BAF normalized to WT, VEH control. Results were compared using a two-way ANOVA followed by a Sidak’s multi-comparisons test. Four asterisks, P<0.0001. n=12. **l**, Scatter dot plot of flow cytometry analysis of the number of LC3-II positive autophagosomes quantified by the mean EGFP-LC3 FI ± s.e.m. from MEFs stably expressing *sgRNA* against *Rosa* (-, black circles) or *Atp7a* (*#1* or *#2*, blue squares, blue triangles) and *EGFP-LC3B* treated with VEH or BAF normalized to *Rosa* (*-*), VEH control. Results were compared using a two-way ANOVA followed by a Sidak’s multi-comparisons test. Three asterisks, P<0.001; Four asterisks, P<0.0001. n=9.

Increased levels of basal autophagy in some cancers represents a vulnerability to autophagy inhibition^5^. BRAF^V600E^-driven lung adenocarcinomas require autophagy to preserve mitochondrial function and supply metabolic substrates to support tumor growth^47^. We previously demonstrated that genetic ablation of *Ctr1* reduced BRAF^V600E^-driven lung tumor development, leading to a survival advantage^4^. While the reliance of BRAF^V600E^-driven tumors on both autophagy^47^ and Cu transport^4^ could be exploited therapeutically, activating mutations in *KRAS* are the most common driver event in this disease^48^. Importantly, oncogenic *Kras^G12D^*-driven lung adenocarcinomas also depend on efficient autophagy for tumor maintenance^49^. Deletion of the critical autophagy component *Atg7* converts *Kras^G12D^*-driven adenomas and adenocarcinomas to oncocytomas, a benign tumor type characterized by an accumulation of dysfunctional mitochondria^49^. To investigate the relationship between Cu, oncogenic KRAS-driven lung cancer, and autophagy, we investigated whether loss of *Ctr1* disrupts autophagy in an autochthonously arising KRAS^G12D^-driven lung adenocarcinoma model. We transduced cohorts of *LSL-**<u>K</u>**ras^G12D/+^*;*Tr**<u>p</u>**53^flox/flox^;Rosa26::LSL-**<u>Luc</u>*** (KPLuc) mice via intratracheal administration of lentivirus expressing Cre recombinase, Cas9, and sgRNA targeting either *β-GAL* or *Ctr1*. Tumor growth was monitored over time with luminescence imaging. Strikingly, we found that genetic targeting of Cu transport blunts the development of KRAS^G12D^-driven lung tumors (Fig. 3a,b and Supplementary Fig. 3a,b). Tumor volume at 16 weeks was significantly decreased in mice with CRISPR-mediated deletion of *Ctr1* (Fig. 3b,d,j). *Ctr1* loss was confirmed by increased CCS levels in the tumor tissue (Fig. 3c,e,k). Molecularly, the established *Ctr1* deficient KPLuc tumors had decreased LC3-positive staining, increased p62-positive staining, and reduced autophagosome number, indicative of diminished autophagy (Fig. 3f,g,h,l,m,n). Thus, the loss of *Ctr1* mitigates KRAS^G12D^-mediated autophagy that is necessary for lung tumor growth.

**Figure 3.**
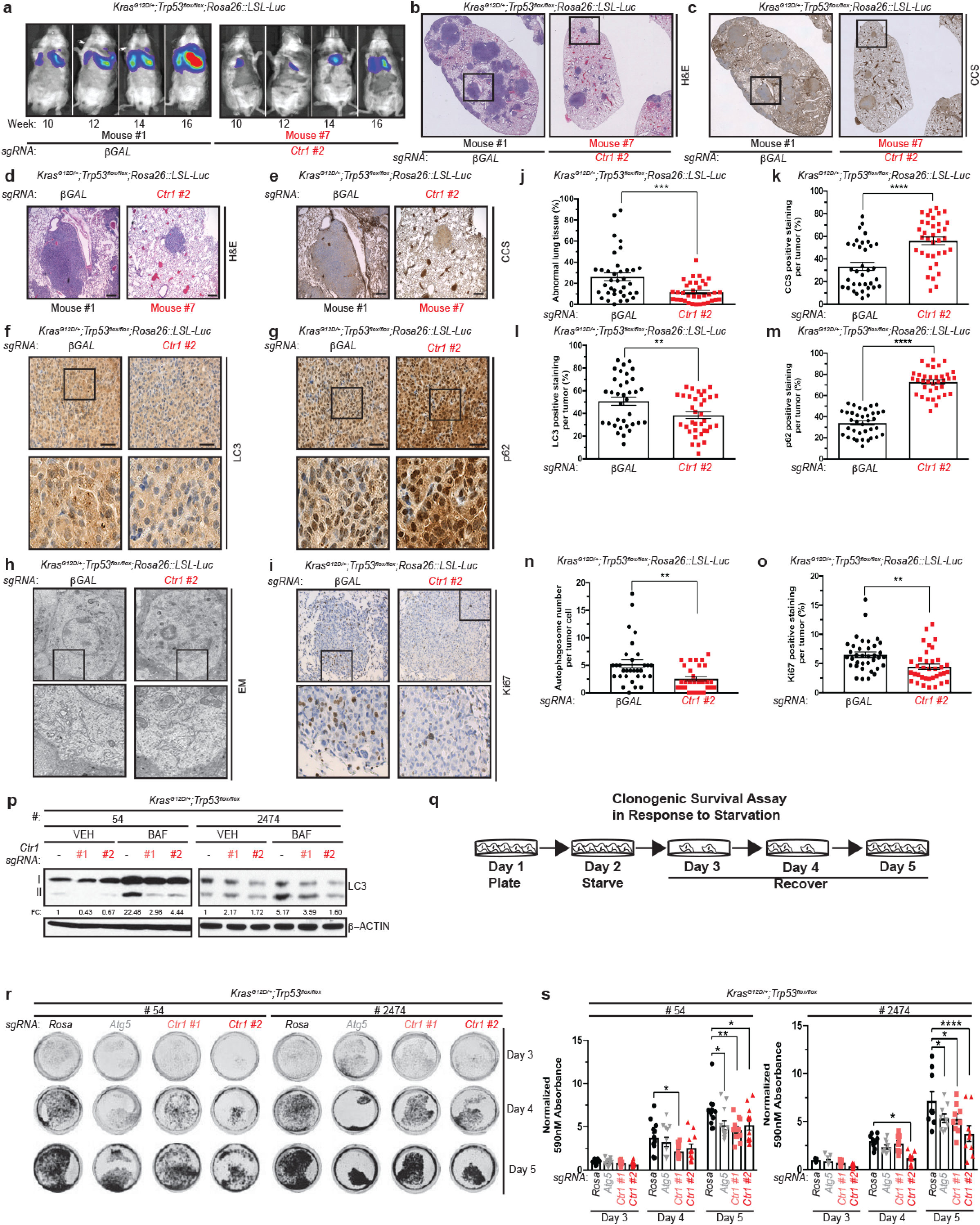
Genetic ablation of *Ctr1* decreases autophagy and proliferation to reduce tumorigenesis and sensitizes cancer cells to starvation in a mouse model of *Kras^G12D^*-driven lung cancer. **a**, Normalized representative images of *in vivo* luminescence of *Kras^G12D/+^*;*Tr<u>p</u>53^flox/flox^;Rosa26::LSL-Luc* (**KPLuc**) mice introduced with either *sgRNA* against *β-GAL* or *Ctr1* at indicated time points. **b**,**c**,**d**,**e**,**f**,**g**,**h**,**i**, Representative images of H&E stained (**b**-5x stitched images, **d**-5x); immunohistochemical detection of CCS (**c**-5x stitched images, **e**-5x), LC3 (**f**-40x), p62 (**g**-40x), or Ki67 (**i**-40x); or electron microscopy (EM) (**h**) of lungs from KPLuc mice expressing *sgRNA* against *β-GAL* or *Ctr1* (5x scale bar: 250 μm; 40x scale bar: 50 μm). **j**,**k**,**l**,**m**,**n**,**o**, Scatter dot plot of mean ± s.e.m. % abnormal lung tissue (**j**, *β-GAL*, n=36 and *Ctr1*, n=35), % CCS positive staining per tumor (**k**, *β-GAL*, n=35 and *Ctr1*, n=36), % LC3 positive staining per tumor (**l**, n=35), % p62 positive staining per tumor (**m**, n=35), autophagosome number per tumor cell (**n**, *β-GAL*, n=31 and *Ctr1*, n=30), or % Ki67 positive staining per tumor (**o**, n=36) from KPLuc mice expressing *sgRNA* against *β-GAL* (black circles) or *Ctr1* (red squares). Results were compared using an unpaired, one-tailed Student’s t-test. Two asterisks, P<0.01; Three asterisks, P<0.001; Four asterisks, P<0.0001. **p**, Immunoblot detection of LC3-I, LC3-II, or β-ACTIN from KP lung adenocarcinoma cell lines #54 (KP #54) and #2474 (KP #2474) stably expressing *sgRNA* against *Rosa* (-) or *Ctr1 (#1* or *#2*) treated with vehicle (VEH) or bafilomycin (BAF). Quantification: ΔLC3-II/β-ACTIN normalized to *Rosa* (*-*), VEH control. **q**, Schematic of clonogenic survival assay in response to starvation. KP #54 and KP #2474 cells stably expressing *sgRNA* against *Rosa, Atg5,* or *Ctr1* following one day of starvation (EBSS) were recovered for three days in normal medium (DMEM). **r**, Representative crystal violet images of KP #54 and KP #2474 cells stably expressing *sgRNA* against *Rosa, Atg5,* or *Ctr1* (*#1* or *#2*) from days 3, 4, and 5 of recovery. **s**, Scatter dot plot of mean absorbance of extracted crystal violet at 590nM ± s.e.m. of KP #54 and KP #2474 cells stably expressing *sgRNA* against *Rosa* (black circles)*, Atg5* (grey inverted triangles), or *Ctr1* (*#1* or *#2,* pink squares, red triangles) from days 3, 4, and 5 of recovery normalized to *Rosa*, day 3 control. Results were compared using a two-way ANOVA followed by a Tukey’s multi-comparisons test. One asterisk, P<0.05; Two asterisks, P<0.01; Four asterisks, P<0.0001. KP#54, n≥11; KP #2474, n=12. Western blot images are representative of at least three biological replicates.

Interestingly, several recent studies reported that genetic disruption of oncogenic KRAS or MAPK pathway inhibition elicits protective autophagy via an ULK1-dependent signaling mechanism and indicate that dual targeting of the canonical MAPK pathway and autophagy may be necessary for anti-tumorigenic activity in KRAS-driven tumors^50–52^. In agreement with our findings that Cu is necessary for MEK1/2^4^ and ULK1/2 kinase activity (Fig. 1d-g and Supplementary Fig. 1e-h), loss of *Ctr1* in KRAS^G12D^ lung tumors diminished ERK1/2 activation (Supplementary Fig. 3c,d), while also blunting compensatory autophagy induction via ULK1/2 (Fig. 3f,g,h,l,m,n). These data suggest that the dual inhibition of these kinase signaling nodes contributes to the reduced *in vivo* tumor cell proliferation (Ki67-positive staining, Fig. 3i,o) associated with *Ctr1* deficiency. However, these results failed to address the extent to which Cu-regulated MAPK signaling versus Cu-mediated autophagy contributes to lung tumor cell growth and survival during KRAS^G12D^-driven tumorigenesis. To begin to interrogate whether the requirement for Cu to induce autophagy is necessary for oncogenic KRAS^G12D^ lung tumor cell phenotypes, we established lung adenocarcinoma cell lines isolated from *LSL-**<u>K</u>**ras^G12D/+^*;*Tr**<u>p</u>**53^flox/flox^* (KP) mice in which *Ctr1* was inactivated via CRISPR/Cas9 (Supplementary Fig. 3e). Targeted disruption of *Ctr1* reduced LC3-II levels in two KP lung adenocarcinoma cell lines as compared to cells transduced with a non-targeting control sgRNA (Fig. 3p and Supplementary Fig. 3f). The dependency of KRAS-driven cancer cell lines on autophagy for growth and survival has been previously evaluated *in vitro* with clonogenic survival assays (Fig. 3q), in which cancer cells are plated in normal media, subsequently starved of amino acids for one day to induce autophagy, and then recovered in normal media for three days, and the ability of the cancer cells to survive is assessed^53^. In agreement with Guo *et* al.^53^, CRISPR/Cas9 knockout of *Atg5* in KP lung adenocarcinoma cells significantly reduced the clonogenic survival of cells post starvation, an effect that was phenocopied by *Ctr1* deletion (Fig. 3r,s). Importantly, expression of ERK2^GOF^ was not sufficient to rescue the decrease in clonogenic survival in the absence *Atg5* or *Ctr1* (Supplementary Fig. 3g,h), which suggests that the survival of these lung cancer cells depends on autophagy in a MAPK-independent fashion. Nevertheless, our collective results demonstrate that the dependence of KRAS^G12D^-driven tumors on MAPK signaling and autophagy events can be inhibited by altering Cu levels, which could be achieved with Cu chelators alone.

To mechanistically interrogate the contribution of Cu-binding to ULK1 signal transduction and autophagy initiation, we next mutated H136, M188, and H197 to alanine, which share conserved primary sequence with the ***C***u-***b***inding ***m***utant (CBM) of MEK1^4^ (Fig. 1a). Compared to wild-type ULK1, the ULK1^CBM^ had reduced *in vitro* ability to bind a Cu-charged resin and phosphorylate ATG13 (Fig. 4a,b). While the lack of ULK^CBM^ *in vitro* kinase activity could suggest a Cu-binding independent finding, we previously demonstrated that introduction of mutations at these conserved residues in MEK5, which is highly similar to MEK1/2 and does not bind Cu, did not alter MEK5 kinase activity^4^. Along these lines, the ULK1^CBM^ immunoprecipitated from *Ulk1/2^-/-^* MEFs deprived of amino acids failed to phosphorylate ATG13 as compared to the exogenous ULK1^WT^ (Fig. 4c). While both ULK1^WT^ and ULK1^CBM^ were efficiently phosphorylated by AMPK at serine 555 and mTORC1 at serine 757 when stably expressed in *Ulk1/2^-/-^* MEFs (Supplementary Fig. 4a), the ULK1^CBM^ exhibited a mobility shift that corresponded to reduced ULK1 autophosphorylation due to defective kinase activity (Supplementary Fig. 4b). Further, despite decreased kinase activity the ULK1^CBM^ retained strong interactions with canonical binding partners ATG13, ATG101, and FIP200, suggesting that the three-dimensional structure of ULK1 is not destabilized upon introduction of mutations into ULK1^CBM^ (Supplementary Fig. 4c). To evaluate the consequence of disrupting ULK1 Cu-binding on autophagic flux, signaling, and autophagosome formation, *Ulk1* and *Ulk2* knockout MEFs were generated via CRISPR/Cas9 and validated to have reduced autophagic flux in response to amino acid deprivation, but as previously reported still retain visible levels of LC3-II^54^ (Supplementary Fig. 4d-f). Only expression of ULK1^WT^ partially rescued LC3-II levels, phosphorylation of ATG13, and autophagosome formation in the context of *Ulk1/2* deficiency, indicating that Cu binding is required to restore autophagic flux downstream of ULK1 (Fig. 4d-j and Supplementary Fig. 4g,h). Finally, consistent with Cu-dependent autophagy acting solely through ULK1/2, we found that loss of *Ctr1* in MEFs deficient in *Ulk1/2* does not further alter autophagic flux (Fig. 4k and Supplementary Fig. 4i), as measured by LC3-II levels. This further bolsters the conclusion that Cu levels mechanistically contribute to autophagy via ULK1/2 and not another Cu-dependent parallel pathway.

**Figure 4.**
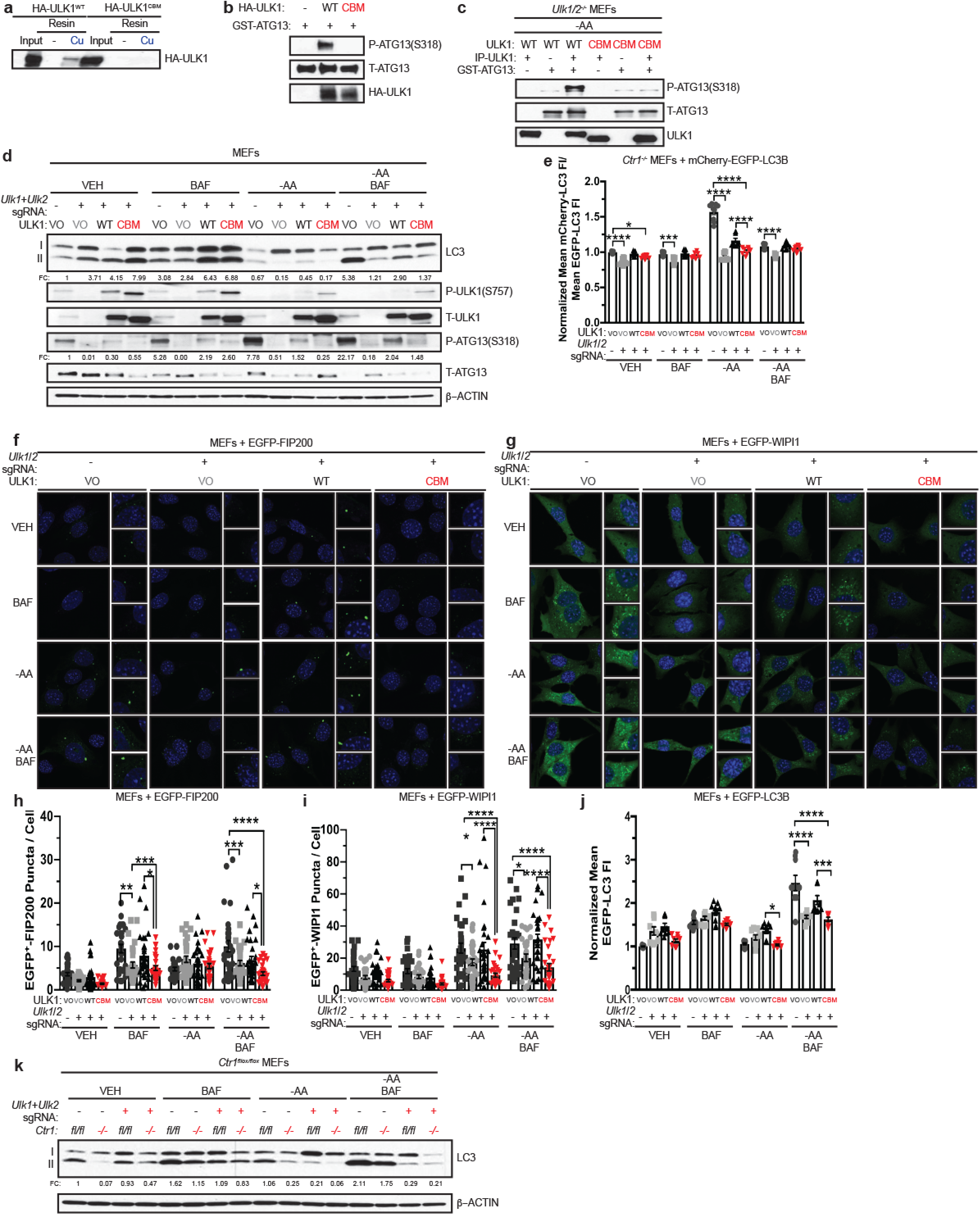
Binding of Cu to ULK1 is required for autophagy signaling and induction. **a**, Immunoblot detection of recombinant HA-ULK1^WT^ or ***C***u ***b***inding ***m***utant HA-ULK1^CBM^ bound to a resin charged with or without Cu compared to input. **b**, Immunoblot detection of recombinant phosphorylated (P)-ATG13, total (T)-ATG13, and HA-ULK1^WT^ or HA-ULK1^CBM^ from ULK1 *in vitro* kinase assays. **c**, Immunoblot detection of recombinant P-ATG13, T-ATG13, or immunoprecipitated (IP)-ULK1 from *Ulk1/2^-/-^* MEFs stably expressing *HA-ULK1^WT^* (WT) or *HA-ULK1^CBM^* (CBM) treated with amino acid deprivation (-AA). **d**, Immunoblot detection of LC3-I, LC3-II, P-ULK1, T-ULK1, P-ATG13, T-ATG13, or β-ACTIN from MEFs stably expressing *sgRNA* against *Rosa* (-) reconstituted with *empty vector* (VO) or *sgRNA* against *Ulk1* and *Ulk2* reconstituted with *empty vector* (VO), *HA-ULK1^WT^* (WT), or *HA-ULK1^CBM^* (CBM) treated with vehicle (VEH) or -AA with or without bafilomycin (BAF). Quantification: ΔLC3-II/β-ACTIN and ΔP-ATG13/T-ATG13 normalized to *Rosa* (-), VO, VEH control. **e**, Scatter dot plot of flow cytometry analysis of autophagic flux quantified by the ratio of mean mCherry-LC3 fluorescent intensity (FI)/mean EGFP-LC3 FI ± s.e.m. from MEFs stably expressing *sgRNA* against *Rosa* (-) reconstituted with VO (dark grey circles) or *sgRNA* against *Ulk1* and *Ulk2* reconstituted with VO (grey squares), WT (black triangles), or CBM (red inverted triangles) and *mCherry-EGFP-LC3B* treated with VEH or -AA with or without BAF normalized to *Rosa* (*-*), VO, VEH control. Results were compared using a two-way ANOVA followed by a Tukey’s multi-comparisons test. One asterisk, P<0.05; Three asterisks, P<0.001; Four asterisks, P<0.0001. n=9. **f**,**g**, Representative fluorescence images of EGFP-FIP200 (**f**) or EGFP-WIP1 (**g**) from MEFs stably expressing *sgRNA* against *Rosa* (-) reconstituted with VO (dark grey circles) or *sgRNA* against *Ulk1* and *Ulk2* reconstituted with VO, WT, or CBM and *EGFP-FIP200* or *EGFP-WIP1* treated with VEH or - AA with or without BAF. **h**,**i**, Scatter dot plot of quantified EGFP^+^ puncta per cell ± s.e.m. from MEFs stably expressing *sgRNA* against *Rosa* (-) reconstituted with VO (dark grey circles) or *sgRNA* against *Ulk1* and *Ulk2* reconstituted with VO (grey squares), WT (black triangles), or CBM (red inverted triangles) and *EGFP-FIP200* (**h**) or *EGFP-WIP1* (**i**) treated with VEH or -AA with or without BAF. Results were compared using a two-way ANOVA followed by a Sidak’s multi-comparisons test. One asterisk, P<0.05; Two asterisks, P<0.01; Three asterisks, P<0.001; Four asterisks, P<0.0001. EGFP-FIP200, n≥26; EGFP-WIP1, n≥27. **j**, Scatter dot plot of flow cytometry analysis of the number of LC3-II positive autophagosomes quantified by the mean EGFP-LC3 FI ± s.e.m. from MEFs stably expressing *sgRNA* against *Rosa* (-) reconstituted with VO (dark grey circles) or *sgRNA* against *Ulk1* and *Ulk2* reconstituted with VO (grey squares), WT (black triangles), or CBM (red inverted triangles) and *EGFP-LC3B* treated with VEH or -AA with or without BAF normalized to *Rosa* (*-*), VO, VEH control. Results were compared using a two-way ANOVA followed by a Sidak’s multi-comparisons test. One asterisk, P<0.05; Three asterisks, P<0.001; Four asterisks, P<0.0001. n=9. **k**, Immunoblot detection of LC3-I, LC3-II, or β-ACTIN from *Ctr1^flox^*^/*flox*^ (*fl/fl*) or *Ctr1^-/-^* (*-/-*) MEFs stably expressing *sgRNA* against *Rosa* (-) or *sgRNA* against *Ulk1* and *Ulk2* (*+*) treated with VEH or -AA with or without BAF. Quantification: ΔLC3-II/β-ACTIN normalized to *Ctr1^flox/flox^*, *Rosa* (-), VEH control. Western blot images are representative of at least three biological replicates.

To translate our finding that Cu-binding to ULK1 is required for autophagy to KRAS-driven tumorigenesis, we next tested and found that blocking ULK1 Cu-binding was associated with reduced tumor growth kinetics and endpoint tumor weight of KRAS^G12D^-transformed *Ulk1^-/-^* immortalized MEFs stably expressing the ULK1^CBM^ compared to ULK1^WT^ (Supplementary Fig. 5a-c). To investigate whether these results could be extrapolated to the setting of endogenous oncogenic KRAS tumorigenesis, *Ulk1* and *Ulk2* were knocked out by CRISPR/Cas9 in mouse *KRAS* mutant lung adenocarcinoma cell lines (Supplementary Fig. 5d). As expected, loss of endogenous *Ulk1/2* decreased the levels of LC3-II and phosphorylation of ATG13 (Fig. 5a,b and Supplementary Fig. 5e,f). Targeted disruption of *Ulk1/2* reduced *in vivo* tumor growth to a similar extent to loss of *Atg5* (Fig. 5c-e). In agreement with Cu being required for ULK1 kinase activity, only ULK1^WT^, but not ULK1^CBM^ could partially restore autophagic flux, signaling, and efficient subcutaneous tumor growth (Fig. 5a-e). Given that Cu mechanistically contributes to autophagy via ULK1/2, the clonogenic survival of KP adenocarcinoma cells was dependent on ULK1/2 but was not further exacerbated by deletion of *Ctr1* (Fig. 5f,g and Supplementary Fig. 5g). Together, these data support a new mechanism to regulate ULK1 activity via Cu binding to promote nutrient deprived or oncogene-driven autophagy that is required for tumorigenesis.

**Figure 5.**
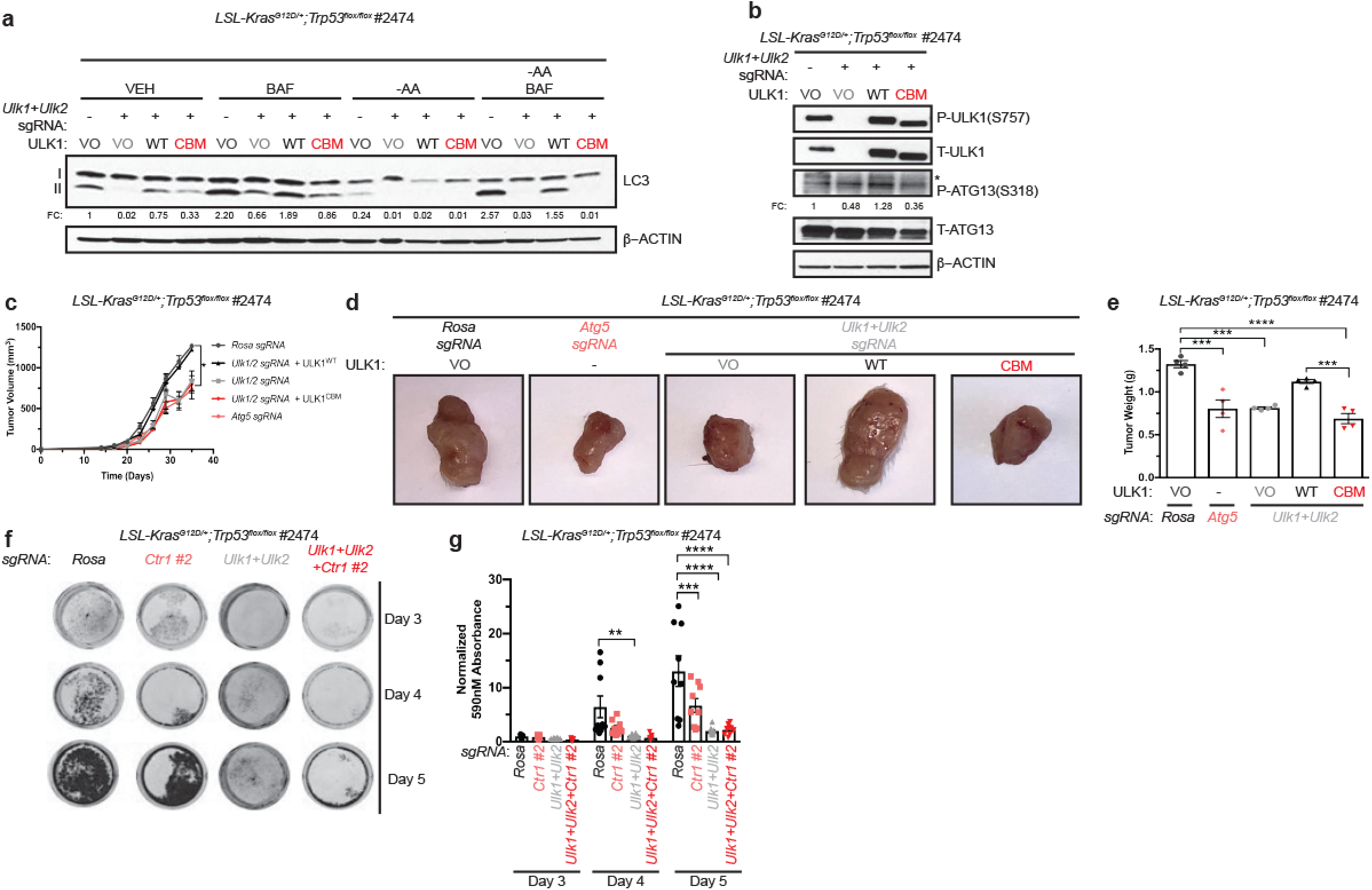
Binding of Cu to ULK1 is required for KRAS^G12D^-driven tumor growth and cancer cell survival in response to starvation. **a**, Immunoblot detection of LC3-I, LC3-II, or β-ACTIN from *Kras^G12D^;p53^flox/flox^* (KP) lung adenocarcinoma cell line #2474 (KP #2474) stably expressing *sgRNA* against *Rosa* (-) reconstituted with *empty vector* (VO) or *sgRNA* against *Ulk1* and *Ulk2* reconstituted with *empty vector* (VO), *HA-ULK1^WT^* (WT), or *HA-ULK1^CBM^* (CBM) treated with vehicle (VEH) or amino acid deprivation (-AA) with or without bafilomycin (BAF). Quantification: ΔLC3-II/β-ACTIN normalized to *Rosa* (-), VO, VEH control. **b**, Immunoblot detection of phosphorylated (P)-ULK1, total (T)-ULK1, P-ATG13, T-ATG13, or β-ACTIN from *Kras^G12D^;p53^flox/flox^* (KP) lung adenocarcinoma cell line #2474 cells stably expressing *sgRNA* against *Rosa* (-) reconstituted with VO or *sgRNA* against *Ulk1* and *Ulk2* reconstituted with VO, WT, or CBM. Quantification: ΔP-ATG13/T-ATG13 normalized to *Rosa* (-), VO, VEH control. **c**, Mean tumor volume (mm^3^) ± s.e.m. versus time (days) in mice injected with KP #2474 cells stably expressing *sgRNA* against *Rosa* reconstituted with VO (dark grey line, dark grey circles)*, Atg5* (pink line, pink hexagons), or *sgRNA* against *Ulk1* and *Ulk2* reconstituted with VO (grey line, grey squares), WT (black line, black triangle), or CBM (red line, red inverted triangle). Results were compared using a paired, two-tailed Student’s t-test. One asterisk, P<0.05. n=4. **d**, Representative dissected tumors from mice injected with KP #2474 cells stably expressing *sgRNA* against *Rosa* (-) reconstituted with VO, *sgRNA* against *Atg5*, or *sgRNA* against *Ulk1* and *Ulk2* reconstituted with VO, WT, or CBM. **e**, Scatter dot plot of mean tumor weight (g) ± s.e.m. of tumors at endpoint from KP #2474 cells stably expressing *sgRNA* against *Rosa* reconstituted with VO (dark grey circles), *Atg5* (pink hexagons), or *sgRNA* against *Ulk1* and *Ulk2* reconstituted with VO (grey squares), WT (black triangle), or CBM (red inverted triangle). Results were compared using a two-way ANOVA followed by a Tukey’s multi-comparisons test. Three asterisks, P<0.001; Four asterisks, P<0.0001. n=4. **f**, Representative crystal violet images of KP #2474 cells stably expressing *sgRNA* against *Rosa*, *Ctr1* (*#2*), *Ulk1 and Ulk2*, *or Ulk1*, *Ulk2*, and *Ctr1 (#2*) from days 3, 4, and 5 of recovery. **g**, Scatter dot plot of mean absorbance of extracted crystal violet at 590nM ± s.e.m. of KP #2474 cells stably expressing *sgRNA* against *Rosa* (black circles)*, Ctr1 (#2,* pink squares), *Ulk1 and Ulk2* (grey triangles), *or Ulk1*, *Ulk2*, and *Ctr1 (#2*, red inverted triangles) from days 3, 4, and 5 of recovery normalized to *Rosa*, day 3 control. Results were compared using a two-way ANOVA followed by a Tukey’s multi-comparisons test. Two asterisks, P<0.01; Three asterisks, P<0.001; Four asterisks, P<0.0001. n≥9. Western blot images are representative of at least three biological replicates.

Despite evidence that Cu homeostasis is essential for life^7^, relatively little is known about cellular pathways that respond directly to Cu levels. We show here that Cu directly modulates the activity of the autophagic kinases ULK1 and ULK2 to function as an endogenous rheostat to control autophagy during nutrient deprivation. These findings highlight a molecular basis for a novel Cu-dependent cellular process to support energy homeostasis. One enticing hypothesis is that sensing Cu abundance through dynamic signaling networks may help cells adapt to environmental changes that would require differential cellular metabolism. By extension, limiting Cu availability or decreasing the binding of Cu to ULK1 may be an attractive therapeutic opportunity to reduce the growth of oncogene-driven lung adenocarcinomas that in part rely on autophagy for tumor maintenance and cell survival. Further, while targeting protein kinase catalytic activity is a proven approach in the landscape of cancer therapeutics, orthogonal approaches that exploit the Cu-dependency of kinases that are critical for maintaining cellular processes essential for tumor growth, such as autophagy, represents an untapped therapeutic opportunity.

## Methods

Methods, including statements of data availability and any associate accession codes and reference are available online.

## Acknowledgements

We thank D.J. Thiele (Duke University), C.J. Chang (University of California Berkeley), R.K. Amaravadi (University of Pennsylvania), and S.A. Tooze (The Francis Crick Institute) for reagents and R.K. Amaravadi, I.A. Asangani, C.V. Dang, J.M. Davis, T.P. Gade, K.E. Hamilton, B. Keith, M.E. Murphy, R. Natesan, A.M. O’Reilly, S.W. Ryeom, M.C. Simon, B.Z. Stanger, J. Tobias, N.A. Tripp, A.T. Weerartna, K.E. Wellen, and E.S. Witze for technical support, discussions, and/or review of the manuscript. This work was supported by American Cancer Society Postdoctoral Fellowship 131203 PF 17 147 01 CCG (J.M.P), NIH grants GM124749 (D.C.B.), CA193603 (D.M.F), CA222503 (D.M.F), ES019851 (M.C), CA243294 (T.T), Pew Scholars Program in Biomedical Science Award #50359 (D.C.B.), and V Foundation Scholar Award 3C59 8ABS 3424 3BDA (D.C.B).

## Author contributions

J.M.P. generated cell lines, performed flow cytometry, immunofluorescence imaging, live cell imaging, immunohistochemical analysis, Western blot analysis, crystal violet growth assays, quantitative PCR (qPCR), and assisted in *in vivo* KRAS^G12D^ mouse tumor model work. T.T. contributed to the study design, generated cell lines and plasmids, prepared samples for ICP-MS, performed Western blot analysis, immunocomplex kinase assays, crystal violet growth assays, co-immunoprecipitations, qPCR, and *in vivo* xenograft mouse work, analyzed data, and prepared figures. A.G. maintained *in vivo* KRAS^G12D^ mouse tumor models and performed *in vivo* KRAS^G12D^ mouse tumor model imaging. M.C. generated tumors *in vivo* KRAS^G12D^ mouse model. D.M.F. contributed to the study design, provided expertise *in vivo* mouse tumor models, and facilitated data analysis. D.C.B conceived of the project, contributed to the study design, generated cell lines, plasmids, and recombinant proteins, performed Cu-binding assays, *in vitro* kinase assays, immunofluorescence imaging, and *in vivo* xenograft mouse work, analyzed data, prepared figures, and wrote the manuscript. All authors read and provided feedback on manuscript and figures.

## Online Methods

### Cell lines

*Ctr1^+/+^* and *Ctr1*^-/-^ immortalized (with SV40) MEFs were previously described and provided by D.J. Thiele (Duke University)^55^. *Ctr1^flox/flox^* MEFs were isolated from homozygous *Ctr1^flox/flox^* embryos and subsequently immortalized *in vitro* via infection with retrovirus expressing SV40^56^. To delete *Ctr1* and establish *Ctr1^-/-^* MEFs, the immortalized *Ctr1^flox/flox^* MEFs were infected *in vitro* with adenovirus expressing Cre-recombinase from pH5’040 CMV PI-Cre (University of Pennsylvania Vector Core). *Atg5^-/-^* MEFs were previously described and provided by R.K. Amaravadi (University of Pennsylvania)^57^. Mouse *LSL-**K**ras^G12D/+^*;*Tr**<u>p</u>**53^flox/flox^* (**KP**) lung adenocarcinoma cell lines were created and cultured as previously described^58^. *Ulk1/2^+/+^*, *Ulk1/2^-/-^*, and *Ulk1^-/-^* immortalized (with SV40) MEFs were previously described and provide by S.A. Tooze (Francis Crick Institute)^54^. *Ctr1^-/-^* immortalized MEFs, KP lung adenocarcinoma cell lines, *Ulk1/2^+/+^* immortalized MEFs, *Ulk1/2^-/-^* immortalized MEFs, and *Ulk1^-/-^* immortalized MEFs were stably infected with retroviruses derived from pBABE or pWZL (*see* plasmids below) or lentiviruses derived from pLentiCRISPRV2 (*see* plasmids below) using established protocols.

### Plasmids

pWZLblasti-CTR1^WT^ (Addgene plasmid #53157)^4^, pBABEpuro-mCherry-EGFP-LC3B (Addgene plasmid #22418)^59^, pBABEpuro-EGFP-LC3B (Addgene plasmid #22405)^60^, pLenti-CRISPRV2blasti (Addgene plasmid #83480), pLenti-CRISPRV2puro (Addgene plasmid #52961)^61^, pWZLblasti-MEK1^WT^ (Addgene plasmid #53161)^4^, pWZL-MEK1^CBM^ (CBM:M187A/H188A/M230A/H239A)^4^, pBABEpuro-HA-ERK2^GOF^ (GOF: R67S/D321N; Addgene plasmid #53203)^4^, pMXs-IP-EGFP-ATG13 (Addgene plasmid #38191)^19^, pMXs-IP-EGFP-FIP200 (Addgene plasmid #38192)^62^, pMXspuro-EGFP-DFCP1 (Addgene plasmid #38269)^39^, pMXs-IP-EGFP-WIPI1 (Addgene plasmid #38272)^39^, and pLentiCRISPRv2Cre (Addgene plasmid #82415)^63^ were previously described. The TRC cloning vector pLKO.1 expressing the mouse *Mek1* shRNA target sequence 5’-CCTGGAGATCAAACCCGCAAT-3’ was obtained from the High-Throughput Screening Core at the University of Pennsylvania. pLentiCRISPRV2puro-*Atg5*-sgRNA expressing the mouse *Atg5* target sequence 5’-TTCCATGAGTTTCCGATTGA was obtained from GenScript. pBABEbleo-mCherry-EGFP-LC3B was created by PCR subcloning mCherry-GFP-LC3B from pBABEpuro-mCherry-EGFP-LC3B plasmid (Addgene plasmid #22418)^59^ with primers designed to N-terminus of mCherry and C-terminus of LC3B. pLenti-CRISPRV2blasti-*Rosa26*-sgRNA was created to express the mouse *Rosa26* target sequence 5’-CCCGATCCCCTACCTAGCCG. pLentiCRISPRV2blasti-*Atp7a*-sgRNA #1 was created to express the mouse *Atp7a* target sequence 5’-TCTATAGGGCAAAACCTCCG. pLentiCRISPRV2blasti-*Atp7a*-sgRNA #2 was created to express the mouse *Atpa7a* target sequence 5’-AGGGATCTTCTACTGCTCTG. pLentiCRISPRV2puro-*Ctr1*-sgRNA #1 was created to express mouse *Ctr1* target sequence 5’-TTGGTAATCAATACACCTGG. pLentiCRISPRV2blasti-*Ctr1*-sgRNA #2 was created to express mouse *Ctr1* target sequence 5’-GGACTCAAGATAGCCCGAGA. pLentiCRISPRV2Cre-*Ctr1*-sgRNA #2 was created to express mouse *Ctr1* target sequence 5’-GGACTCAAGATAGCCCGAGA. pETDUET-6XHIS-TEV-HA-ULK1^WT^, pBABEbleo-HA-ULK1^WT^, and pWZLblasti-HA-ULK1^WT^ were created by PCR subcloning ULK1^WT^ from pRK5-HA-ULK1^WT^ (Addgene plasmid #31963)^20^ with primers designed to include an N-terminal HA-tag. pETDUET-6XHIS-TEV-HA-ULK1^CBM^ (CBM: H136A/M188A/H197A), pBABEbleo-HA-ULK1^CBM^, and pWZLblasti-HA-ULK1^CBM^ were created by introducing mutations corresponding to the indicated amino acid changes by site-directed mutagenesis into ULK1^WT^ from the pWZLblasi-HA-ULK1^WT^ plasmid followed by PCR subcloning with primers designed to include an N-terminal HA-tag for the pETDUET plasmid. pLentiCRISPRV2blasti-*Ulk1*-sgRNA and pLentiCRISPRV2hygro-*Ulk1*-sgRNA were created to express the mouse *Ulk1* target sequence 5’-GGCAGTGTACGGTTCCGAGG. pLentiCRISPRV2blasti-*Ulk2*-sgRNA and pLentiCRISPRV2puro-*Ulk2*-sgRNA were created to express the mouse *Ulk2* target sequence 5’-AGGCCCATGACGAGTAACCA. pBABEbleo-HA-KRAS^G12D^ was created by PCR subcloning KRAS^G12D^ from the pBABEpuro-HA-KRAS^G12D^ (Addgene plasmid #58902) with primers designed to include an N-terminal HA-tag.

### *In vitro* copper binding

100 ng or 1 µg of recombinant GST-ULK1, GST-ULK2, HA-ULK1, or HA-ULK1^CBM^ were incubated in 500 µl of RIPA containing 25 µl Profinity IMAC resin (Bio-Rad) charged with no metal, Cu^2+^, Zn^2+^, or Fe^3+^ for 1 hour at 4 °C. SDS-PAGE analysis and immunoblot were performed as described below.

### Immunoblot analysis

Indicated cell lines were washed with cold PBS and lysed with cold RIPA buffer containing 1X EDTA-free Halt^TM^ protease and phosphatase inhibitor cocktail halt protease and phosphatase inhibitors (Thermo Scientific). The protein concentration was determined by BCA Protein Assay (Pierce) using BSA as a standard. Equal amount of lysates were resolved by SDS-PAGE using standard techniques, and protein was detected with the following primary antibodies: mouse anti-GST (1:5000; 2624, Cell Signaling), rabbit anti-phospho(S318)-ATG13 (1:1000; 600-401-C49, Rockland), rabbit anti-ATG13 (1:1000; 13273, Cell Signaling), rabbit anti-ULK1 (1:1000; 8054, Cell Signaling), rabbit anti-ULK2 (1:1000; NBP1-33136, Novus Biologicals), rabbit anti-phospho(Ser2481)-mTOR (1:1000; 2974, Cell Signaling), rabbit anti-phospho(Ser2448)-mTOR (1:1000; 2971, Cell Signaling), rabbit anti-mTOR (1:1000; 2972, Cell Signaling), mouse anti-phospho(Thr389)-p70S6K (1:1000; 9206, Cell Signaling), rabbit anti-p70S6K (1:1000; 2708, Cell Signaling), rabbit anti-phospho(Ser757)-ULK1 (1:1000; 6888, Cell Signaling), rabbit anti-CCS (1:1000; 20141, Santa Cruz), mouse anti-b-ACTIN (1:10000; 3700, Cell Signaling), rabbit anti-LC3B (1:2000; NB100-2220, Novus Biologicals), rabbit anti-phospho(Thr202/Tyr204)-ERK1/2 (1:1000; 9101, Cell Signaling), mouse anti-ERK1/2 (1:1000; 9107, Cell Signaling), mouse anti-HA (1:1000; 2367, Cell Signaling), rabbit anti-ATG101 (1:1000; 13492, Cell Signaling), or rabbit anti-FIP200 (1:1000; 12436, Cell Signaling) followed by detection with one of the horseradish peroxidase conjugated secondary antibodies: mouse anti-rabbit IgG light chain specific (1:5000; 93702, Cell Signaling), goat anti-rabbit IgG (1:5000, 7074, Cell Signaling), or goat anti-mouse IgG (1:5000, 7076, Cell Signaling), using SignalFire (Cell Signaling) or SignalFire Elite ECL (Cell Signaling) detection reagents. The fold change in the ratio of phosphorylated protein to total protein was measured in Image Studio Lite (LI-CORE Biosciences) software by boxing each band per representative image using the rectangular selection tool, and calculating the total area of the band in pixels. The total area of the phosphorylated protein band in pixels was normalized to the total area of the total protein band in pixels. The average fold change is shown in figures.

### *In vitro* kinase assays

ULK1 and ULK2 *in vitro* kinase assays were performed as previously described for MEK1 kinase assays^3^. Briefly, 100 ng of recombinant GST-ATG13 (Rockland) with or without 1 µg of recombinant GST-ULK1 (Thermo Fisher), GST-ULK2 (Thermo Fisher), HA-ULK1^WT^ (*see* protein purification), or HA-ULK1^CBM^ (*see* protein purification) were incubated in 90 µl of kinase buffer (25 mM Tris-HCl [pH 7.5], 20 mM MgCl2, 2 mM dithiothreitol [DTT], 25 mM β-GP, 0.5 mM Na_3_VO_4_, 120 μM ATP) in the presence or absence of increasing concentrations of CuCl_2_ from 0 to 10 µM, increasing concentrations of TTM (Sigma) from 0 to 50 µM, or 10 µM of MRT68921 at 30 °C for 30 minutes.

### Immunocomplex kinase assays

Immunocomplex ULK1 kinase assays were performed using ULK1 immunoprecipitated from cell lysates. Briefly, indicated cell lines were washed in cold PBS and lysed in cold NP-40 buffer containing 1X EDTA-free Halt^TM^ protease and phosphatase inhibitor cocktail halt protease and phosphatase inhibitors (Thermo Scientific). The protein concentration was determined by BCA Protein Assay (Pierce) using BSA as a standard. 1.5mg of total protein lysate was used to immunoprecipitate endogenous or exogenous ULK1 with rabbit anti-ULK1 antibody (1:50; 8054, Cell Signaling) at 4°C overnight, followed by incubation with 20 µl Protein A agarose bead slurry (Cell Signaling) for 3 hours. Beads were washed three times with NP-40 buffer, once with PBS, then resuspended in 90 µl kinase buffer (see *in vitro* kinase assays). 100-500 ng of recombinant GST-ATG13 (Rockland) was added to each reaction and incubated at 30°C for 30 minutes.

### Flow cytometry

Indicated cell lines stably expressing pBABEbleo-mCherry-EGFP-LC3B, pBABEpuro-mCherry-EGFP-LC3B, or pBABEpurp-EGFP-LC3B were washed with PBS and collected for flow analysis as previously described^37, 46^. Briefly, 80,000 cells/well were plated on 6-well plates and 48 hours later treated with vehicle, 500 nM rapamycin (Selleck Chemicals), or EBSS media (Sigma) with or without 100 nM bafilomycin (Cayman) for 3 hours or increasing concentrations of tetrathiomolybdate (TTM, Sigma) or deferoxamine mesylate salt (DFO, Sigma) for 24 hours. Cells were harvested with trypsin, spun down at 2000 rpm for 4 minutes, and then resuspended in PBS without (autophagic flux) or with (autophagosome number) 0.1% saponin twice. Samples were run on an Attune NxT flow cytometer in triplicate and 30,000 events were captured per sample. Data analysis was carried out with FlowJo 8.7. Statistical analysis of autophagic flux (ratio of normalized mean mCherry-LC3 fluorescence intensity/mean EGFP-LC3 fluorescence intensity) or autophagosome number (normalized mean EGFP-LC3 fluorescence intensity) was analyzed using a two-way ANOVA followed by a Sidak’s multi-comparisons test or a two-way ANOVA followed by a Tukey’s multi-comparisons test in Prism 7 (GraphPad).

### Immunofluorescence

Indicated cell lines stably expressing pBABEpuro-mCherry-EGFP-LC3B were plated on glass microslides and 48 hours later treated with vehicle, 100 nM bafilomycin, 500 nM rapamycin, or the media was changed to EBSS for 3 hours, washed with PBS, fixed in 4% paraformaldehyde in 1XPBS for 10 minutes, washed PBS five times, and mounted immediately on slides with ProLong Gold Antifade Mountant with DAPI (Thermo Fisher). Indicated cell lines stably expressing pMXs-IP-EGFP-ATG13, pMXs-IP-EGFP-FIP200, pMXspuro-EGFP-DFCP1, or pMXs-IP-EGFP-WIPI1 were plated on glass microslides and 48 hours later treated with regular media with or without 100 nM bafilomycin or EBSS media with or without 100 nM bafilomycin for 3 hours, washed with PBS, fixed in 4% paraformaldehyde in 1XPBS for 10 minutes, washed PBS five times, and mounted immediately on slides with ProLong Gold Antifade Mountant with DAPI (Thermo Fisher). Slides were imaged on a Zeiss LSM 880 inverted confocal imaging system with a 63x oil objective at 2x zoom. Quantification of mCherry-LC3 positive puncta and mCherry EGFP-LC3 positive puncta was performed using the Red and Green Puncta Colocalization Macro created by NIH funded Daniel D.J. Shiwarski (University of Pittsburgh), R.K. Dagda, (University of Nevada), C.T. Chu (University of Pittsburgh) in Image J. DAPI positive cells were manually counted per image. Statistical analysis of mCherry^+^LC3 puncta per cell or mCherry^+^ EGFP^+^LC3 puncta per cell was analyzed using a two-way ANOVA followed by a Sidak’s multi-comparisons test in Prism 7 (GraphPad). Quantification of EGFP positive puncta was performed in ImageJ by splitting the image channel and either utilizing the auto-threshold function RenyiEntropy or manually thresholding for puncta followed by particle analysis set to measure size (pixel^2^) from 5-infinity. DAPI positive cells were manually counted per image. Statistical analysis of mCherry_+_LC3 puncta per cell, mCherry_+_ EGFP_+_LC3 puncta per cell, or EGFP positive puncta per cell was analyzed using a two-way ANOVA followed by a Sidak’s multi-comparisons test in Prism 7 (GraphPad).

### Live cell imaging

For time course imaging of Cu levels, MEFs were seeded at 250,000 cells/plate in 35mm glass insert imaging plates (MatTek) and 24 hours later, 1 μM of CF4^31^ or Ctrl-CF4^31^ was added to the media. After 20 minutes, cells were washed with PBS and either the media was replaced with Live Cell Imaging Solution (Thermo Scientific) containing 1 μM of CF4 or Ctrl-CF4 and treated with vehicle or the media was changed to HBSS (Corning) containing 1 μM of CF4 or Ctrl-CF4. Cells were then imaged on a Zeiss LSM880 laser scanning confocal microscopy system with a 20x dry objective for one hour acquiring images every 5 minutes. Cells were excited at 458nm and images acquired for transmitted light and emission at 521nm. Fluorescence of CF4 or Ctrl-CF4 from single cells was analyzed 10 cells per field of view from three independent images. ImageJ image analysis software was utilized to set thresholds for background, the freehand selection tool was used to select an individual cell, and the measure tool was used to measure the mean intensity, area, and standard deviation. Statistical analysis of normalized CF4 or Ctrl-CF4 fluorescence per cell was performed using a two-way ANOVA followed by a Tukey’s multi-comparisons test in Prism 7 (GraphPad). For Cu levels in *Ctr1^+/+^*, *Ctr1^-/-^*, or CTR1^WT^ reconstituted *Ctr1^-/-^* MEFs, cell lines were seeded at 150,000 cells/plate in 35mm glass insert imaging plates (MatTek) and 48 hours later were washed with PBS and media was replaced with Live Cell Imaging Solution (Thermo Scientific) containing 1 μM of CF4^31^ or Ctrl-CF4^31^ for 20 minutes and imaged on a Zeiss LSM880 laser scanning confocal microscopy system with a 20x dry objective. Cell were excited at 458nm using 8% laser power and images acquired for transmitted light and emission at 521nm. Fluorescence of CF4 or Ctrl-CF4 from single cells was analyzed from three independent experiments in which 10 cells per field of view from three independent images. The total of 90 cells were graphed as a spread with each point representing one cell. ImageJ image analysis software was utilized to set thresholds for background, the freehand selection tool was used to select an individual cell, and the measure tool was used to measure the mean intensity, area, and standard deviation. Statistical analysis of normalized CF4 or Ctrl-CF4 fluorescence per cell was performed using a two-way ANOVA followed by a Sidak’s muti-comparisons test or a one-way ANOVA followed by a Tukey’s multi-comparisons test in Prism 7 (GraphPad).

### Mouse lung cancer model

*LSL-KRAS^G12D^*;*Trp53^flox/flox^; Rosa26::LSL-Luc* mice were maintained on a mixed C567B16/129Sv4 background and were previously described^64, 65^. Mice were given lentivirus expressing Cas9, Cre recombinase, and *β-Gal* sgRNA or *Ctr1* sgRNA from pLentiCRISPRv2Cre generated as previously described at 6x10^4^ pfu per mouse by intratracheal intubation at 6-10 weeks of age as previously described^63^. For *in vivo* bioluminescence imaging, mice were anesthetized and injected with D-Luciferin (GoldBio LUCNA-1G) in PBS at 300 mg/kg. Luminescent signals were acquired 20 minutes post injection with the IVIS Spectrum (Caliper Life Sciences). Analysis was done using Living Image 4.5 (Perkin Elmer). All studies were approved by the University of Pennsylvania Institutional Animal Care and Use Committee.

### Immunohistochemistry

Sections were deparaffinized, rehydrated and subjected to epitope retrieval stained with a rabbit anti-LC3B antibody (1:1000; NB100-2220, Novus Biologicals), rabbit anti-p62 antibody (1:1000; PM04, MBL International), mouse anti-CCS (1:100; 55561, Santa Cruz Biotechnology), rabbit anti-Ki67 (1:1000; VP-RM04, Vector Laboratories), or rabbit anti-phospho(Thr202/Tyr204)-ERK1/2 (1:1000; 4370, Cell Signaling) followed by peroxidase-based detection and counterstaining with haematoxylin using the Leica Bond Rx^m^ system. Photographs were taken on a Leica DMI6000B inverted light and fluorescent microscope (20x for quantitation, 5x or 40x for representative images). LC3-, p62-, and CCS-positive staining as well as tumor burden was assessed in Image J. The color deconvolution macro was applied to images and tumors were circumscribed with the freehand selection tool on the DAB staining (Color_2) generated window. Using the threshold function, the total area of the tumor in pixels was recorded utilizing the same parameters for each tumor image. Areas staining positive by these parameters were selected and the positive-staining area in pixels was recorded. The positive-staining area of the tumor in pixels was divided by the total area of the tumor in pixels to determine the percentage positive-staining area. Statistical analysis of abnormal lung area, % positive CCS, % positive LC3, % positive p62, or % P-ERK1/2 staining was analyzed using an unpaired, two-tailed Student’s t-test in Prism 7 (GraphPad).

### Electron Microscopy

Tumor tissue was processed and prepared for ultrastructure analysis by the Electron Microscopy Resource Lab (EMRL) at the University of Pennsylvania. Specimens were imaged on a JEOL JEM 1010 transmission electron microscope equipped with a 2k x 2k AMT CCD camera which uses a tungsten filament as its electron source. Autophagosome number per cell were manually counted by a researcher blinded to the treatment group. Statistical analysis of autophagosome number was analyzed using an unpaired, two-tailed Student’s t-test in Prism 7 (GraphPad).

### Clonogenic survival assays

Cells were seeded at 80,000 cells/well in 12-well plates and 24 hours later washed with PBS and media was changed to EBSS. After 24 hours, EBSS was removed and replaced with media. Cells were fixed and stained with 500uL crystal violet solution (0.5%) for 15 minutes after 24, 48, and 72 hours. Cells were washed with deionized water and plates allowed to dry before images were taken with the BioRad Chemidoc Imaging System. Cells were destained by adding 1 mL of 10% acetic acid/well and plates were rocked for 20 minutes at room temperature. Samples from each well were transferred to a 96-well plate, and 590nM absorbance was read on a Synergy HT plate reader (BioTek Instruments). Statistical analysis of 590nM absorbance was performed using a two-way ANOVA followed by Tukey’s multi-comparison test in Prism 7 (Graphpad).

### Protein purification

Recombinant GST-ULK1 (Thermo Scientific), GST-ULK2 (Thermo Scientific), and GST-ATG13 (Rockland) were purchased. HIS-TEV-HA-ULK1^WT^ and HIS-TEV-HA-ULK1^CBM^ were expressed from in BL21-CodonPlus(DE3)-RIL competent cells (Agilent Technologies) in Terrific Broth (Fisher Scientific) at 37 °C until reaching an OD600 of 0.6-1.0 and then induced with 1 mM IPTG overnight at 18 °C. Bacteria cultures were ultracentrifuged at 8,000 r.p.m and resuspended in lysis buffer (25 mM HEPES pH 7.5, 500 mM NaCl, 10% glycerol, 1 mM PMSF, 10 mM 2-Mercaptoethanol, complete EDTA free protease inhibitor cocktail [Pierce]). Lysate was sonicated at 50% duty cycle for 30 second pulses two times on ice or 50% duty cycle for 30 pulses on and 30 pulses off for 30 minutes at 4°C. Lysate was centrifuged at 12,000 r.p.m. for 15 minutes at 4°C. Supernatant was incubated with Ni Sepharose 6 Fast Flow beads (GE Healthcare) at 4°C rocking for 3 hours and subsequently eluted via gravity column and the resin was then washed with 1 L of wash buffer (25 mM HEPES pH 7.5, 500 mM NaCl, 10% glycerol, 40 mM Imidazole, 10 mM 2-Mercaptoethanol). The protein was then eluted with with 25 mM HEPES pH 7.5, 500 mM NaCl, 10% glycerol, 300 mM Imidazole, 10 mM 2-Mercaptoethanol and the elution fractions were pooled together and treated with TEV protease overnight while dialyzing into dialysis buffer 1 (25 mM HEPES pH 7.5, 50 mM NaCl, 10% glycerol, 10 mM 2-Mercaptoethanol). The eluent was then applied to a HiTrap Q HP anion exchange 5mL column and eluted over a gradient of 20 column volumes ranging from 0% to 100% of Buffer B (25 mM HEPES pH 7.5, 1 M NaCl). The protein elutes between 15% and 30% Buffer B. Peak fractions were run on an SDS-PAGE gel, pooled, concentrated, and run on a Superdex S200 10/300 gel filtration column in a final buffer of 25 mM HEPES pH7.5, 150 mM NaCl, 5% Glycerol, 10 mM 2-Mercaptoethanol. Protein was concentrated, flash frozen in liquid nitrogen, and stored in -80 °C freezer for future use.

### Mouse xenografts

10^7^ KP lung adenocarcinoma cells or immortalized, transformed MEFs resuspended in phosphate buffer saline were injected subcutaneously into flanks of SCID/beige mice (Strain 250, Charles River Laboratory) as previously described^66^. All studies were approved by the University of Pennsylvania Institutional Animal Care and Use Committee. Statistical analysis of tumor volumes and tumor weights at end point was performed using an unpaired, one-tailed Student’s t-test in Prism 7 (GraphPad).

### Reverse transcriptase-PCR and Reverse transcriptase-quantitative PCR

For RT-PCR, RNA was purified from MEFs and reverse transcribed to cDNA using standard protocols and then PCR amplified with the primers 5’-CTGTTTTCCGGTTTGGTGAT-3’ and 5’-TGCCCAACAGTTTTGTGTGT-3’ to detect human *CTR1 or* 5’-GCACAGTCAAGGCCGAGAAT-3’ and 5’-GCCTTCTCCATGGTG GTGAA-3’ to detect mouse *Gapdh*. For RT-qPCR, RNA was purified from MEFs or tumor cell lines and reverse transcribed to cDNA as previously described and then quantified utilizing a Taqman probe Mm00437663_m1 to detect mouse *Atp7a*, Mm00558247_m1 to detect mouse *Ctr1*, Mm00437238_m1 to detect mouse *Ulk1*, Mm03048846_m1 to detect mouse *Ulk2*, or Mm01277042_m1 to detect mouse TATA-binding protein (*Tbp*) using the ViiA 7 Real-Time PCR System. Relative mRNA expression levels were normalized to *Tbp* and analyzed using comparative delta-delta CT method.

### Inductively coupled mass spectrometry

Indicated cell lines were washed with PBS and cell pellet was harvested by centrifugation. Inductively coupled mass spectrometry of Cu parts per million (ppm) levels was analyzed at University of Pennsylvania School of Veterinary Medicine. Statistical analysis of Cu (ppm) per weight of tissue was performed using a one-way ANOVA followed by a Tukey’s multi-comparisons test in Prism 7 (GraphPad).

### Statistics and reproducibility

Data are represented as the mean ± s.e.m. The sample size (*n*) indicates the experimental replicates, cells, or mice analyzed. For all experiments, each data point analyzed was from an independent biological sample. Statistical significance was typically determined using an unpaired, one-tailed Student’s t-test, an unpaired, two-tailed Student’s t-test, a one-way ANOVA followed by a Tukey’s multi-comparisons test, a two-way ANOVA followed by a Sidak’s multi-comparisons test, or a two-way ANOVA followed by Tukey’s multi-comparisons test, in which significance was determined as P≤ 0.05.

### Reporting Summary

Further information on experimental design is available in the Nature Research Reporting Summary linked to this article.

## References

1. Fleuren, E. D. G., Zhang, L., Wu, J. & Daly, R. J. The kinome ‘at large’ in cancer. Nat. Rev. Cancer 16, 83– 98 (2016).

2. Rubino, J. T. & Franz, K. J. Coordination chemistry of copper proteins: how nature handles a toxic cargo for essential function. J. Inorg. Biochem. 107, 129–143 (2012).

3. Turski, M. L. et al. A novel role for copper in Ras/MAPK signaling. Mol. Cell. Biol. 32, 1284–1295 (2012).

4. Brady, D. C. et al. Copper is required for oncogenic BRAF signalling and tumorigenesis. Nature 509, 496– 496 (2014).

5. Amaravadi, R. K., Kimmelman, A. C. & Debnath, J. Targeting Autophagy in Cancer: Recent Advances and Future Directions. Cancer Discov. 9, 1167–1181 (2019).

6. Kolch, W., Halasz, M., Granovskaya, M. & Kholodenko, B. N. The dynamic control of signal transduction networks in cancer cells. Nat. Rev. Cancer 15, 515–527 (2015).

7. Festa, R. A. & Thiele, D. J. Copper: an essential metal in biology. Curr. Biol. 21, R877–83 (2011).

8. Chelly, J. et al. Isolation of a candidate gene for Menkes disease that encodes a potential heavy metal binding protein. Nat. Genet. 3, 14–19 (1993).

9. Mercer, J. F. B. et al. Isolation of a partial candidate gene for Menkes disease by positional cloning. Nat. Genet. 3, 20–25 (1993).

10. Huster, D. et al. Consequences of copper accumulation in the livers of the Atp7b-/-(Wilson disease gene) knockout mice. Am. J. Pathol. 168, 423–434 (2006).

11. Pfeiffenberger, J. et al. Hepatobiliary malignancies in Wilson disease. Liver Int. 35, 1615–1622 (2015).

12. Krishnamoorthy, L. et al. Copper regulates cyclic-AMP-dependent lipolysis. Nat. Chem. Biol. 12, 586–592 (2016).

13. Brady, D. C., Crowe, M. S., Greenberg, D. N. & Counter, C. M. Copper chelation inhibits BRAFV600E-driven melanomagenesis and counters resistance to BRAFV600E and MEK1/2 inhibitors. Cancer Res. 77, 6240–6252 (2017).

14. Xu, M. M., Casio, M., Range, D. E., Sosa, J. A. & Counter, C. M. Copper chelation as targeted therapy in a mouse model of oncogenic BRAF-driven papillary thyroid cancer. Clin. Cancer Res. 24, 4271–4281 (2018).

15. Goodman, V. L., Brewer, G. J. & Merajver, S. D. Control of copper status for cancer therapy. Curr. Cancer Drug Targets 5, 543–549 (2005).

16. Petherick, K. J. et al. Pharmacological inhibition of ULK1 kinase blocks mammalian target of rapamycin (mTOR)-dependent autophagy. J. Biol. Chem. 290, 11376–11383 (2015).

17. Chan, E. Y. W., Kir, S. & Tooze, S. A. siRNA Screening of the Kinome Identifies ULK1 as a Multidomain Modulator of Autophagy. J Biol Chem 282, 25464–25474 (2007).

18. Ganley, I. G. et al. ULK1.ATG13.FIP200 complex mediates mTOR signaling and is essential for autophagy. J Biol Chem 284, 12297–12305 (2009).

19. Hosokawa, N. et al. Nutrient-dependent mTORCl association with the ULK1-Atg13-FIP200 complex required for autophagy. Mol. Biol. Cell 20, 1981–1991 (2009).

20. Jung, C. H. et al. ULK-Atg13-FIP200 complexes mediate mTOR signaling to the autophagy machinery. Mol Biol Cell 20, 1992–2003 (2009).

21. Kim, D.-H. et al. mTOR Interacts with Raptor to Form a Nutrient-Sensitive Complex that Signals to the Cell Growth Machinery. Cell 110, 163–175 (2002).

22. Thoreen, C. C. et al. A unifying model for mTORC1-mediated regulation of mRNA translation. Nature 485, 109–113 (2012).

23. Hara, K. et al. Amino Acid Sufficiency and mTOR Regulate p70 S6 Kinase and eIF-4E BP1 through a Common Effector Mechanism. J Biol Chem 273, 14484–14494 (1998).

24. Kim, E., Goraksha-Hicks, P., Li, L., Neufeld, T. P. & Guan, K.-L. Regulation of TORC1 by Rag GTPases in nutrient response. Nat Cell Biol 10, 935–945 (2008).

25. Sancak, Y. et al. Ragulator-Rag complex targets mTORC1 to the lysosomal surface and is necessary for its activation by amino acids. Cell 141, 290–303 (2010).

26. Inoki, K., Zhu, T. & Guan, K.-L. TSC2 Mediates Cellular Energy Response to Control Cell Growth and Survival. Cell 115, 577–590 (2003).

27. Gwinn, D. M. et al. AMPK phosphorylation of raptor mediates a metabolic checkpoint. Mol Cell 30, 214– 226 (2008).

28. Sabatini, D. M., Erdjument-Bromage, H., Lui, M., Tempst, P. & Snyder, S. H. RAFT1: A mammalian protein that binds to FKBP12 in a rapamycin-dependent fashion and is homologous to yeast TORs. Cell 78, 35–43 (1994).

29. Brown, E. J. et al. A mammalian protein targeted by G1-arresting rapamycin–receptor complex. Nature 369, 756–758 (1994).

30. Brady, G. F. et al. Regulation of the copper chaperone CCS by XIAP-mediated ubiquitination. Mol Cell Biol 30, 1923–1936 (2010).

31. Xiao, T. et al. Copper regulates rest-activity cycles through the locus coeruleus-norepinephrine system. Nat. Chem. Biol. 14, 655–663 (2018).

32. Chiang, G. G. & Abraham, R. T. Phosphorylation of Mammalian Target of Rapamycin (mTOR) at Ser-2448 Is Mediated by p70S6 Kinase. J Biol Chem 280, 25485–25490 (2005).

33. Brown, E. J. et al. Control of p70 s6 kinase by kinase activity of FRAP in vivo. Nature 377, 441–446 (1995).

34. Soliman, G. A., et al. mTOR Ser-2481 Autophosphorylation Monitors mTORC-specific Catalytic Activity and Clarifies Rapamycin Mechanism of Action. J Biol Chem 285, 7866–7879 (2010).

35. Kim, J., Kundu, M., Viollet, B. & Guan, K.-L. AMPK and mTOR regulate autophagy through direct phosphorylation of Ulk1. Nat Cell Biol 13, 132–141 (2011).

36. Ichimura, Y. et al. A ubiquitin-like system mediates protein lipidation. Nature 408, 488–492 (2000).

37. Gump, J. M. et al. Autophagy variation within a cell population determines cell fate through selective degradation of Fap-1. Nat Cell Biol 16, 47–54 (2014).

38. Pankiv, S. et al. p62/SQSTM1 binds directly to Atg8/LC3 to facilitate degradation of ubiquitinated protein aggregates by autophagy. J. Biol. Chem. 282, 24131–24145 (2007).

39. Itakura, E. & Mizushima, N. Characterization of autophagosome formation site by a hierarchical analysis of mammalian Atg proteins. Autophagy 6, 764–776 (2010).

40. Mercer, T. J., Gubas, A. & Tooze, S. A. A molecular perspective of mammalian autophagosome biogenesis. J Biol Chem 293, 5386–5395 (2018).

41. Karanasios, E. et al. Dynamic association of the ULK1 complex with omegasomes during autophagy induction. J Cell Sci 126, 5224–5238 (2013).

42. Egan, D. F. et al. Small Molecule Inhibition of the Autophagy Kinase ULK1 and Identification of ULK1 Substrates. Mol. Cell 59, 285–297 (2015).

43. Russell, R. C. et al. ULK1 induces autophagy by phosphorylating Beclin-1 and activating VPS34 lipid kinase. Nat Cell Biol 15, 741–750 (2013).

44. Park, Y. S. et al. AKT-induced PKM2 phosphorylation signals for IGF-1-stimulated cancer cell growth. Oncotarget 7, 48155–48167 (2016).

45. Dooley, H. C. et al. WIPI2 links LC3 conjugation with PI3P, autophagosome formation, and pathogen clearance by recruiting Atg12-5-16L1. Mol Cell 55, 238–252 (2014).

46. Eng, K. E., Panas, M. D., Hedestam, G. B. K. & McInerney, G. M. A novel quantitative flow cytometry-based assay for autophagy. Autophagy 6, 634–641 (2010).

47. Strohecker, A. M. et al. Autophagy Sustains Mitochondrial Glutamine Metabolism and Growth of BrafV600E–Driven Lung Tumors. Cancer Discov. 3, 1272–1285 (2013).

48. Cancer Genome Atlas Research Network. Comprehensive molecular profiling of lung adenocarcinoma. Nature 511, 543–550 (2014).

49. Guo, J. Y. et al. Autophagy suppresses progression of K-ras-induced lung tumors to oncocytomas and maintains lipid homeostasis. Genes Dev. 27, 1447–1461 (2013).

50. Bryant, K. L. et al. Combination of ERK and autophagy inhibition as a treatment approach for pancreatic cancer. Nat Med 25, 628–640 (2019).

51. Kinsey, C. G. et al. Protective autophagy elicited by RAF→MEK→ERK inhibition suggests a treatment strategy for RAS-driven cancers. Nat Med 25, 620–627 (2019).

52. Lee, C.-S. et al. MAP kinase and autophagy pathways cooperate to maintain RAS mutant cancer cell survival. Proc. Natl. Acad. Sci. U. S. A. 116, 4508–4517 (2019).

53. Guo, J. Y. et al. Autophagy provides metabolic substrates to maintain energy charge and nucleotide pools in Ras-driven lung cancer cells. Genes Dev. 30, 1704–1717 (2016).

54. McAlpine, F., Williamson, L. E., Tooze, S. A. & Chan, E. Y. W. Regulation of nutrient-sensitive autophagy by uncoordinated 51-like kinases 1 and 2. Autophagy 9, 361–373 (2013).

## References

55. Lee, J., Peña, M. M. O., Nose, Y. & Thiele, D. J. Biochemical characterization of the human copper transporter Ctr1. J Biol Chem 277, 4380–4387 (2002).

56. Nose, Y., Kim, B.-E. & Thiele, D. J. Ctr1 drives intestinal copper absorption and is essential for growth, iron metabolism, and neonatal cardiac function. Cell Metab. 4, 235–244 (2006).

57. Mizushima, N. et al. Dissection of autophagosome formation using Apg5-deficient mouse embryonic stem cells. J. Cell Biol. 152, 657–668 (2001).

58. Walter, D. M. et al. RB constrains lineage fidelity and multiple stages of tumour progression and metastasis. Nature 568, 423–427 (2019).

59. N’Diaye, E. N. et al. PLIC proteins or ubiquilins regulate autophagy-dependent cell survival during nutrient starvation. EMBO Rep. 10, 173–179 (2009).

60. Fung, C., Lock, R., Gao, S., Salas, E. & Debnath, J. Induction of autophagy during extracellular matrix detachment promotes cell survival. Mol. Biol. Cell 19, 797–806 (2008).

61. Sanjana, N. E., Shalem, O. & Zhang, F. Improved vectors and genome-wide libraries for CRISPR screening. Nat. Methods 11, 783–784 (2014).

62. Hara, T. et al. FIP200, a ULK-interacting protein, is required for autophagosome formation in mammalian cells. J Cell Biol 181, 497–510 (2008).

63. Walter, D. M. et al. Systematic In Vivo Inactivation of Chromatin-Regulating Enzymes Identifies Setd2 as a Potent Tumor Suppressor in Lung Adenocarcinoma. Cancer Res. 77, 1719–1729 (2017).

64. Jackson, E. L. et al. The differential effects of mutant p53 alleles on advanced murine lung cancer. Cancer Res 65, 10280–10288 (2005).

65. Safran, M. et al. Mouse Reporter Strain for Noninvasive Bioluminescent Imaging of Cells that have Undergone Cre-Mediated Recombination: *Mol*. Imaging 2, 297–302 (2003).

66. Hamad, N. M. et al. Distinct requirements for Ras oncogenesis in human versus mouse cells. Genes Dev. 16, 2045–2057 (2002).

