## Supplemental Figures and Legends for "Copper is an essential regulator of the autophagic kinases ULK1/2 to drive lung adenocarcinoma"

### Tsang Posimo *et al.* Supplemental Figure 1

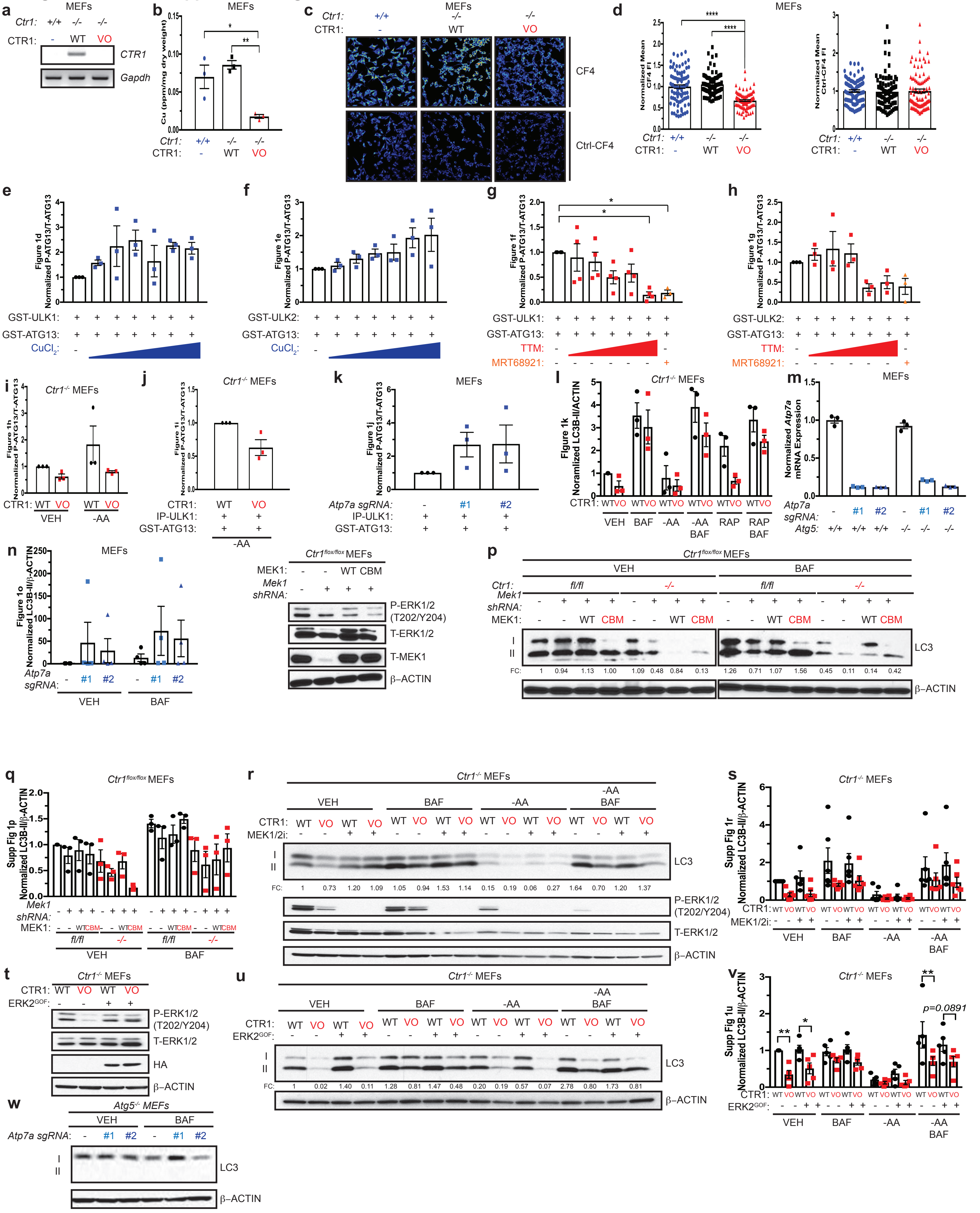

### Tsang Posimo *et* al. Supplemental Figure 2

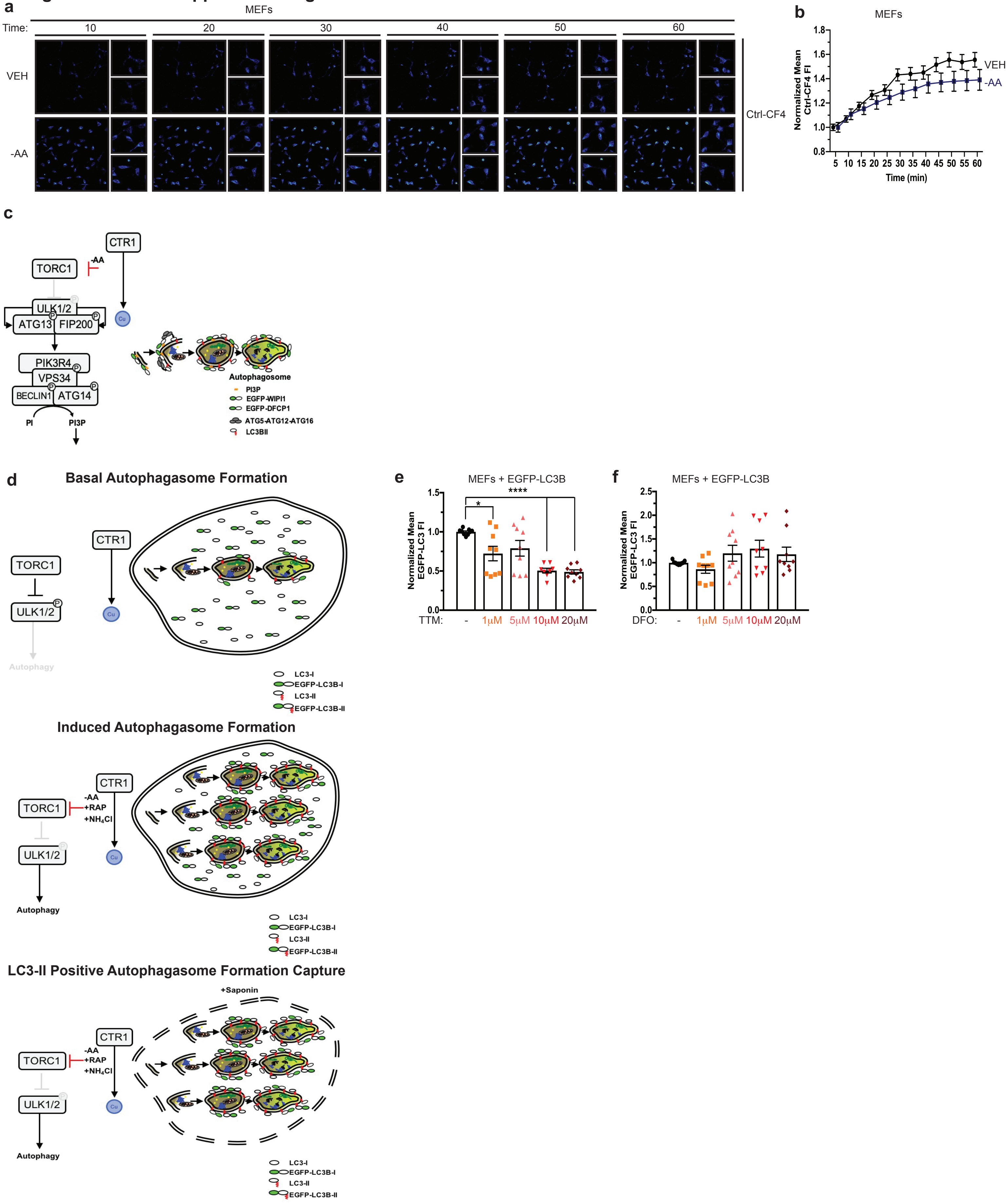

**Tsang Posimo *et* al. Supplemental Figure 3**

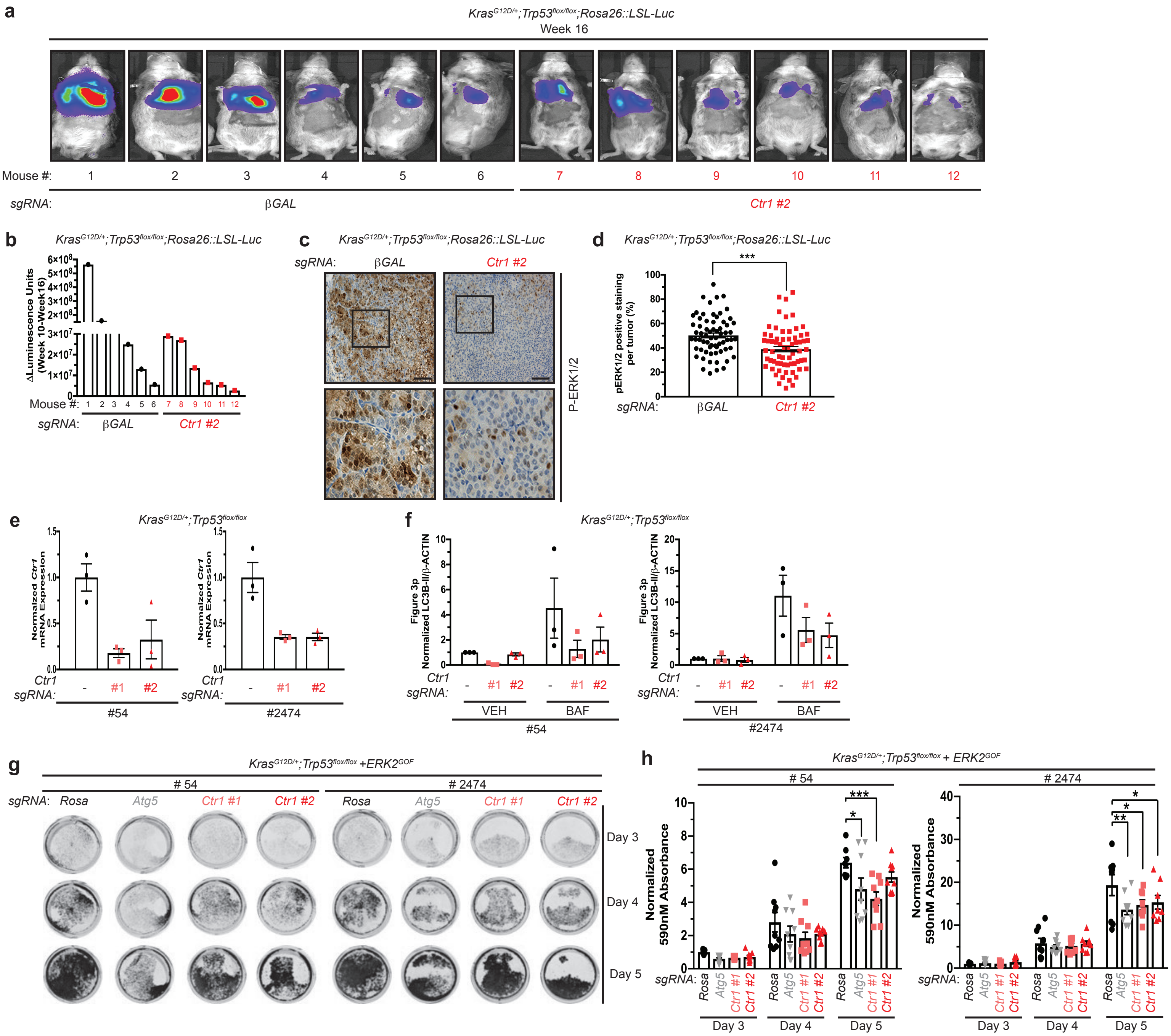

#### Tsang Posimo *et* al. Supplemental Figure 4

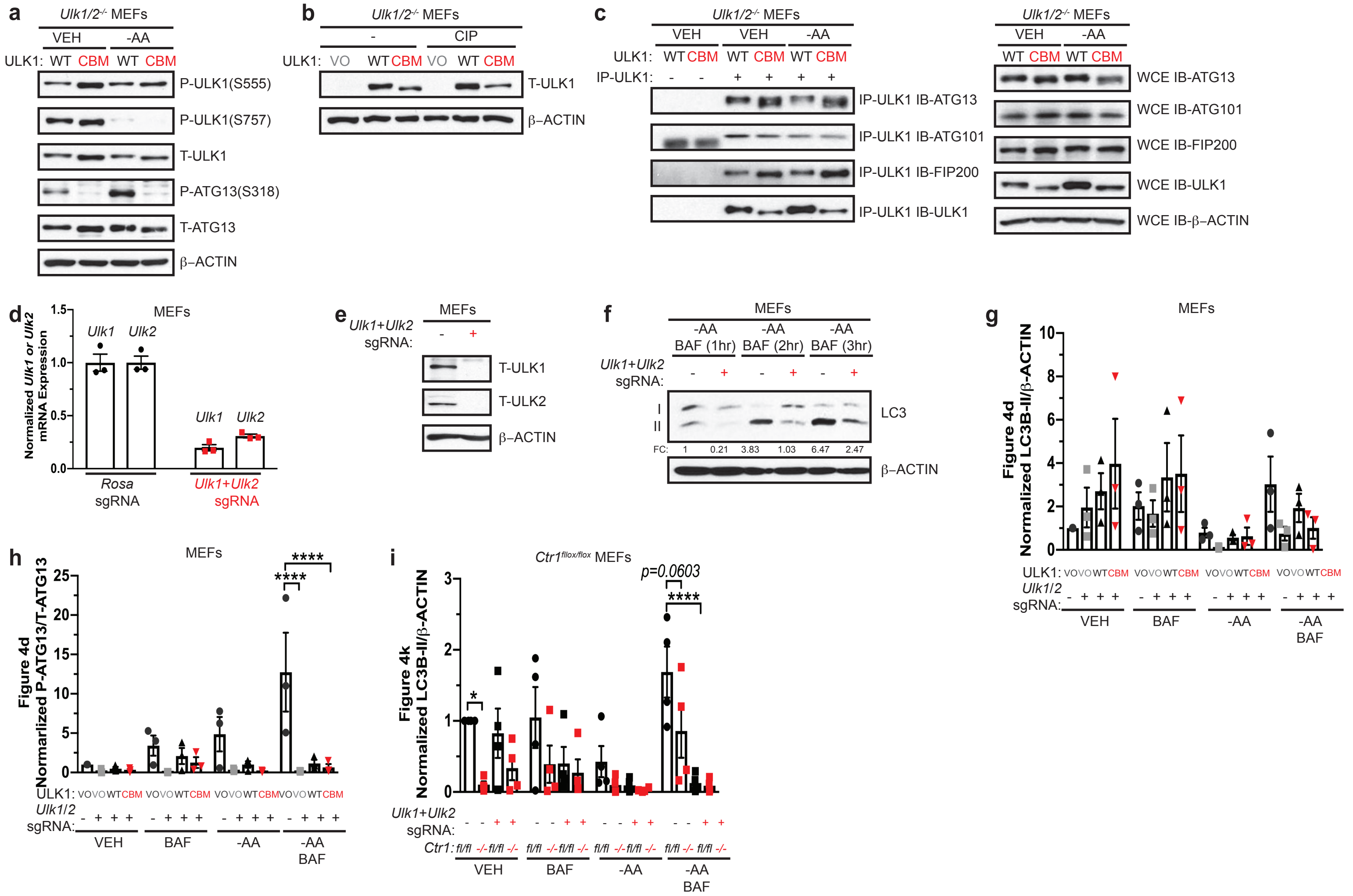

#### Tsang Posimo *et al.* Supplemental Figure 5

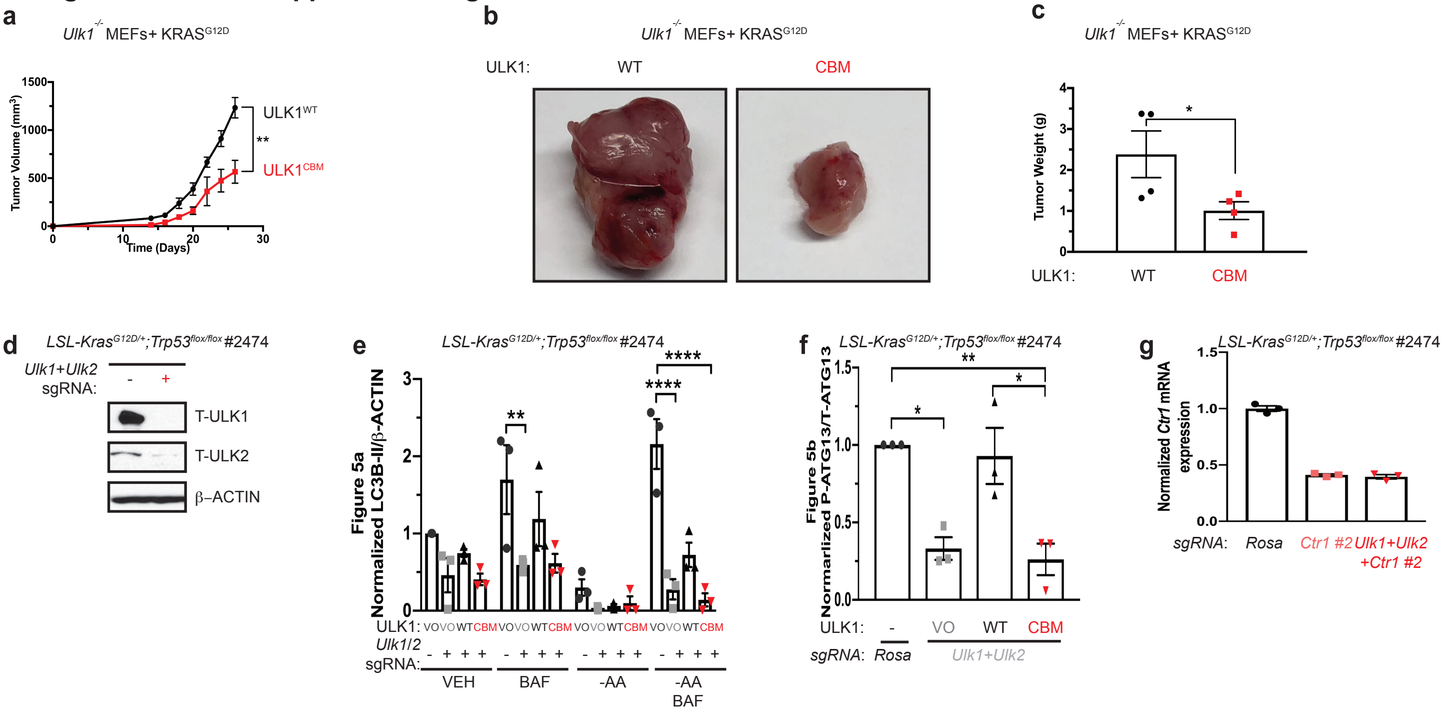

#### Supplementary Figure Legends

**Supplementary Figure 1. Cu is both necessary and sufficient for autophagy induction and signaling in a MAPK signaling independent fashion, upstream of ATG5. a**, RT-PCR detection of indicated mRNAs from *Ctr1*$^{+/+}$ MEFs or *Ctr1*$^{-/-}$ MEFs stably expressing *CTR1*$^{WT}$ (WT) or *empty vector* (VO). **b**, Scatter dot plot of inductively coupled plasma mass spectrometry (ICP-MS) detection of mean Cu (parts per million, ppm) from *Ctr1*$^{+/+}$ MEFs (+/+, blue circles) or *Ctr1*$^{-/-}$ MEFs stably expressing WT (black squares) or VO (red triangles) per sample weight ± s.e.m. Results were compared using a one-way ANOVA followed by a Tukey's multi-comparisons test. One asterisk, P<0.05; Two asterisks, P<0.01. n=3. **c**, Representative live cell imaging of the Cu probe CF4 or control Cu probe Ctrl-CF4 from *Ctr1*$^{+/+}$ MEFs (+/+) or *Ctr1*$^{-/-}$ MEFs stably expressing WT or VO. **d**, Scatter dot plot of quantified mean CF4 or Ctrl-CF4 fluorescence intensity (FI) ± s.e.m. from *Ctr1*$^{+/+}$ MEFs (+/+, blue circles) or *Ctr1*$^{-/-}$ MEFs stably expressing WT (black squares) or VO (red triangles). Results were compared using a one-way ANOVA followed by a Tukey's multi-comparisons test. Four asterisks, P<0.0001. n=90 individual cells. **e**,**f**,**g**,**h**,**i**,**j**,**k**,**l**, Scatter dot plot of quantified normalized ΔPhosphorylated (P)-ATG13/Total (T)-ATG13 or ΔLC3-II/β-ACTIN from **Figure 1d**,**e**,**f**,**g**,**h**,**i**,**j**,**k**. Results were compared using a one-way ANOVA or two-way ANOVA followed by a Tukey's multi-comparisons test. One asterisk, P<0.05. n=3 or 4. **m**, Scatter dot plot of normalized quantitative PCR (qPCR) expression of *Atp7a* mRNA from MEFs or *Atg5*$^{-/-}$ MEFs stably expressing *sgRNA* against *Rosa* (-, black circles) or *Atp7a* (*#1*, blue squares, or *#2*, blue triangles). **n**, Scatter dot plot of quantified normalized ΔLC3-II/β-ACTIN from **Figure 1o**. Results were compared using a two-way ANOVA followed by a Tukey's multi-comparisons test. ns. n=4. **o**, Immunoblot detection of P-ERK1/2, T-ERK1/2, T-MEK1, or β-ACTIN from *Ctr1*$^{flox/flox}$ MEFs stably expressing *empty vector* (-) or *shRNA* against *Mek1* reconstituted with *HA-MEK1*$^{WT}$ (WT) or *HA-MEK1*$^{CBM}$ (CBM). **p**, Immunoblot detection of LC3-I, LC3-II, or β-ACTIN from *Ctr1*$^{flox/flox}$ (*fl/fl*) MEFs or *Ctr1*$^{-/-}$ (-/-) MEFs stably expressing empty vector (-) or *shRNA* against *Mek1* reconstituted with WT or CBM treated with vehicle (VEH) or bafilomycin (BAF). Quantification: ΔLC3-II/β-ACTIN normalized to *fl/fl*, empty vector (-), VEH control. **q**, Scatter dot plot of quantified normalized ΔLC3-II/β-ACTIN from **Supplementary Figure 1p**. Results were compared using a two-way ANOVA followed by a Tukey's multi-comparisons test. ns. n=3. **r**, Immunoblot detection of LC3-I, LC3-II, P-ERK1/2, T-ERK1/2, or β-ACTIN from *Ctr1*$^{-/-}$ MEFs stably expressing WT or VO treated with VEH or amino acid deprivation (-AA) with or without BAF in the presence or absence of MEK1/2 inhibitor (MEK1/2i). Quantification: ΔLC3-II/β-ACTIN normalized to WT, VEH control. **s**, Scatter dot plot of quantified normalized ΔLC3-II/β-ACTIN from **Supplementary Figure 1r**. Results were compared using a two-way ANOVA followed by a Tukey's multi-comparisons test. ns. n=6. **t**, Immunoblot detection of P-ERK1/2, T-ERK1/2, HA, or β-ACTIN from *Ctr1*$^{-/-}$ MEFs stably expressing WT or VO with or without ***g***ain-***o***f-***f***unction (GOF) *HA-ERK2* (ERK2$^{GOF}$). **u**, Immunoblot detection of LC3-I, LC3-II, or β-ACTIN from *Ctr1*$^{-/-}$ MEFs stably expressing WT or VO with or without ERK2$^{GOF}$ with VEH or -AA with or without BAF. Quantification: ΔLC3-II/β-ACTIN normalized to WT, VEH control. **v**, Scatter dot plot of quantified normalized ΔLC3-II/β-ACTIN from **Supplementary Figure 1u**. Results were compared using a two-way ANOVA followed by a Tukey's multi-comparisons test. One asterisk, P<0.05; Two asterisks, P<0.01. n=5. **w**, Immunoblot detection of LC3-I, LC3-II, or β-ACTIN from *Atg5*$^{-/-}$ MEFs stably expressing *sgRNA* against *Rosa* (-) or *Atp7a* (#1 or #2) treated with VEH or BAF. Western blot images are representative of at least three biological replicates.

**Supplementary Figure 2. Cu but not Fe is required for autophagosome formation and is associated with fluctuations in the Cu labile pool**. **a**, Representative live cell imaging of the Cu probe Ctrl-CF4 every ten minutes for 60 minutes from MEFs treated with vehicle (VEH) or amino acid deprivation (-AA). **b**, Quantification of mean Ctrl-CF4 fluorescence intensity (FI) ± s.e.m. versus time (minutes, min) from MEFs treated with VEH (black circles) or -AA (blue squares) normalized to t=0, five minutes. Results were compared using a two-way ANOVA followed by a Sidak's multi-comparisons test. n=30. **c**, Schematic of immunofluorescence-based approach to access autophagosome formation. **d**, Schematic of flow cytometry-based approach to access autophagosome number. **e**,**f**, Scatter dot plot of flow cytometry analysis of the number of LC3-II positive autophagosomes quantified by the mean GFP-LC3 fluorescent intensity ± s.e.m. from MEFs stably expressing *EGFP-LC3B* treated VEH or increasing concentration of Cu chelator TTM (**e**) or Fe chelator DFO (**f**). Results were compared using a one-way ANOVA followed by a Dunnett's multi-comparisons test. One asterisk, P<0.05; Four asterisks, P<0.0001. n=9.

**Supplementary Figure 3. Genetic ablation of *Ctr1* decreases MAPK signaling to reduce *Kras*$^{G12D}$-driven lung tumorigenesis, while survival in response starvation is independent of**

**MAPK signaling.** **a**, Normalized representative images of *in vivo* luminescence of ***K****ras*$^{G12D/+}$;*Tr**p**53*$^{flox/flox}$*;Rosa26::LSL-**Luc*** (**KPLuc**) mice introduced with either *sgRNA* against *β-GAL* or *Ctr1* at week 16 endpoint. **b**, ΔLuminescence units from week 10 to week 16 from KPLuc mice introduced with either *sgRNA* against *β-GAL* (black circles) or *Ctr1* (red squares). **c**, Representative 40x images of immunohistochemical detection of phosphorylated (P)-ERK1/2 of lungs from KPLuc mice expressing *sgRNA* against *β-GAL* or *Ctr1*. (40x scale bar: 50 μm). **d**, Scatter dot plot of mean ± s.e.m. % P-ERK1/2 positive staining per tumor (*β-GAL* and *Ctr1*, n=66) from KPLuc mice expressing *sgRNA* against *β-GAL* (black circles) or *Ctr1* (red squares). Results were compared using an unpaired, one-tailed Student's t-test. Three asterisks, P<0.001. **e**, Scatter dot plot of normalized quantitative PCR (qPCR) expression of *Ctr1* mRNA from KP lung adenocarcinoma cell lines #54 (KP #54) and #2474 (KP #2474) stably expressing *sgRNA* against *Rosa* (-, black circles) or *Ctr1* (*#1*, pink squares or *#2*, red triangles). **f**, Scatter dot plot of quantified normalized ΔLC3-II/β-ACTIN from **Figure 3p**. Results were compared using a two-way ANOVA followed by a Tukey's multi-comparisons test. ns. n=3. **g**, Representative crystal violet images of KP #54 and KP #2474 cells stably expressing *sgRNA* against *Rosa, Atg5,* or *Ctr1* (*#1* or *#2*) and ERK2$^{GOF}$ from days 3, 4, and 5 of recovery. **h**, Scatter dot plot of mean absorbance of extracted crystal violet at 590nM ± s.e.m. of KP #54 and KP #2474 cells stably expressing *sgRNA* against *Rosa* (blue circles)*, Atg5* (grey inverted triangles)*,* or *Ctr1* (*#1* or *#2,* pink squares, red triangles) and ***g**ain-**o**f-**f**unction* (GOF) *HA-ERK2* (ERK2$^{GOF}$) from days 3, 4, and 5 of recovery normalized to *Rosa*, day 3 control. Results were compared using a two-way ANOVA followed by a Tukey's multi-comparisons test. One asterisk, P<0.05; Two asterisks, P<0.01; Three asterisks, P<0.001. KPLuc #54, n≥8; KPLuc #2474, n≥8.

**Supplementary Figure 4. Binding of Cu to ULK1 is required for kinase activity but not substrate association or phosphorylation.** **a**, Immunoblot detection of phosphorylated (P) ATG13, total (T)- ATG13, P-ULK1 (S555), P-ULK1 (S757), T-ULK1, or β-ACTIN from *Ulk1/2*$^{-/-}$ MEFs stably expressing *HA-ULK1*$^{WT}$ (WT) or *HA-ULK1*$^{CBM}$ (CBM) treated with vehicle (VEH) or amino acid deprivation (-AA). **b**, Immunoblot detection of T-ULK1 or β-ACTIN from cell lysates treated with or without calf alkaline phosphatase (CIP) from *Ulk1/2*$^{-/-}$ MEFs stably expressing *empty vector* (VO), *HA-ULK1*$^{WT}$ (WT), or *HA-ULK1*$^{CBM}$ (CBM). **c**, Immunoblot detection of T-ATG13, T-ATG101, T-FIP200, T-ULK1, or β-ACTIN from immunoprecipitated

(IP)-ULK1 or whole cell extracts (WCE) from *Ulk1/2*$^{-/-}$ MEFs stably expressing *HA-ULK1*$^{WT}$ (WT) or *HA-ULK1*$^{CBM}$ (CBM) treated with VEH or -AA. **d**, Scatter dot plot of normalized quantitative PCR (qPCR) expression of *Ulk1* or *Ulk2* mRNA from MEFs stably expressing *sgRNA* against *Rosa* (black circles) or *Ulk1* and *Ulk2* (red squares). **e**, Immunoblot detection of T-ULK1, T-ULK2, or β-ACTIN from MEFs stably expressing *sgRNA* against *Rosa* (-) or *Ulk1* and *Ulk2* (+). **f**, Immunoblot detection of LC3-I, LC3-II, or β-ACTIN from MEFs stably expressing *sgRNA* against *Rosa* (-) or *Ulk1* and *Ulk2* (+) treated with -AA and bafilomycin (BAF) for 1 hour (hr), 2 hr, and 3hr. **g**,**h**,**i**, Scatter dot plot of quantified normalized ΔLC3-II/β-ACTIN or ΔP-ATG13/T-ATG13 from **Figure 4d**,**k**. Results were compared using a two-way ANOVA followed by a Tukey's multi-comparisons test. Four asterisks, $P<0.0001$. n=3.

**Supplementary Figure 5. Binding of Cu to ULK1 is required for tumorigenesis by oncogenic KRAS$^{G12D}$. a**, Mean tumor volume (mm$^3$) ± s.e.m. versus time (days) in mice injected with *Ulk1*$^{-/-}$ MEFs stably expressing either *HA-ULK1*$^{WT}$ or *HA-ULK1*$^{CBM}$ and transformed with KRAS$^{G12D}$. Results were compared using a paired, one-tailed Student's t-test. Two asterisks, $P<0.01$. n=4. **b**, Representative dissected tumors from mice injected with *Ulk1*$^{-/-}$ MEFs stably expressing either *HA-ULK1*$^{WT}$ (WT) or *HA-ULK1*$^{CBM}$ (CBM) and transformed with KRAS$^{G12D}$. **c**, Scatter dot plot of mean tumor weight (g) ± s.e.m. of tumors at endpoint from *Ulk1*$^{-/-}$ MEFs stably expressing either WT (black circles) or CBM (red squares) and transformed with KRAS$^{G12D}$. Results were compared using an unpaired, one-tailed Student's t-test. One asterisk, $P<0.05$. n=4. **d**, Immunoblot detection of T-ULK1, T-ULK2, or β-ACTIN from ***Kras***$^{G12D/+}$;*Trp53*$^{flox/flox}$ (KP) lung adenocarcinoma cell line #2474 (KP #2474) stably expressing *sgRNA* against *Rosa* (-) or *Ulk1* and *Ulk2* (+). **e**,**f**, Scatter dot plot of quantified normalized ΔLC3-II/β-ACTIN or ΔP-ATG13/T-ATG13 from **Figure 5a**,**b**. Results were compared using a two-way ANOVA followed by a Tukey's multi-comparisons test. One asterisk, $P<0.05$; Two asterisks, $P<0.01$; Four asterisks, $P<0.0001$. n=3. **g**, Scatter dot plot of normalized quantitative PCR (qPCR) expression of *Ctr1* mRNA from KP #2474 cells stably expressing *sgRNA* against *Rosa* (black circles), *Ctr1 (#2*, pink squares), or *Ulk1*, *Ulk2*, and *Ctr1 (#2*, red inverted triangles).

**Supplementary Video 1.** Representative live cell imaging of the Cu probe CF4 every ten minutes for 60 minutes from MEFs treated with vehicle (VEH).

**Supplementary Video 2.** Representative live cell imaging of the Cu probe CF4 every ten minutes for 60 minutes from MEFs treated with amino acid deprivation (-AA).

**Supplementary Video 3.** Representative live cell imaging of the Cu probe Ctrl-CF4 every ten minutes for 60 minutes from MEFs treated with vehicle (VEH).

**Supplementary Video 4.** Representative live cell imaging of the Cu probe Ctrl-CF4 every ten minutes for 60 minutes from MEFs treated with amino acid deprivation (-AA).
